# BenzoHTag, a fluorogenic self-labeling protein developed using molecular evolution

**DOI:** 10.1101/2023.10.29.564634

**Authors:** Bryan J. Lampkin, Benjamin J. Goldberg, Joshua A. Kritzer

## Abstract

Self-labeling proteins are powerful tools in chemical biology as they enable the precise cellular localization of a synthetic molecule, often a fluorescent dye, with the genetic specificity of a protein fusion. HaloTag7 is the most popular self-labeling protein due to its fast labeling kinetics and the simplicity of its chloroalkane ligand. Reaction rates of HaloTag7 with different chloroalkane-containing substrates is highly variable and rates are only very fast for rhodamine-based dyes. This is a major limitation for the HaloTag system because fast labeling rates are critical for live-cell assays. Here, we report a molecular evolution system for HaloTag using yeast surface display that enables the screening of libraries up to 10^8^ variants to improve reaction rates with any substrate of interest. We applied this method to produce a HaloTag variant, BenzoHTag, which has improved performance with a fluorogenic benzothiadiazole dye. The resulting system has improved brightness and conjugation kinetics, allowing for robust, no-wash fluorescent labeling in live cells. The new BenzoHTag-benzothiadiazole system has improved performance in live-cell assays compared to the existing HaloTag7-silicon rhodamine system, including saturation of intracellular enzyme in under 100 seconds and robust labeling at dye concentrations as low as 7 nM. It was also found to be orthogonal to the silicon HaloTag7-rhodamine system, enabling multiplexed no-wash labeling in live cells. The BenzoHTag system, and the ability to optimize HaloTag for a broader collection of substrates using molecular evolution, will be very useful for the development of cell-based assays for chemical biology and drug development.

## Introduction

Genetically encoded sensors have become essential in the biochemical sciences.^1, 2^ Such biosensors were originally developed using fluorescent proteins, but advancements in both synthetic dye chemistry and protein engineering have enabled improved chemo-genetic constructs for cellular imaging. Specifically, self-labeling proteins such as HaloTag7 allow organic dyes, which are often brighter and more photostable than fluorescent proteins, to be introduced to cellular environments with chemical and genetic precision.^3-9^ HaloTag7 reacts with a linear chloroalkane ligand which can be appended to any molecule of interest, most commonly fluorescent dyes (Fig. 1a) but also a wide variety of substrates including biomolecules. HaloTag7 has been used in a wide array of cell-based and live-animal assays that have revealed new insights into protein-protein interactions, drug action at the cell surface, cell-surface receptor recycling, targeted protein degradation, and cell penetration of peptide and oligonucleotide drugs, among other areas.^10-15^

**Figure 1.**
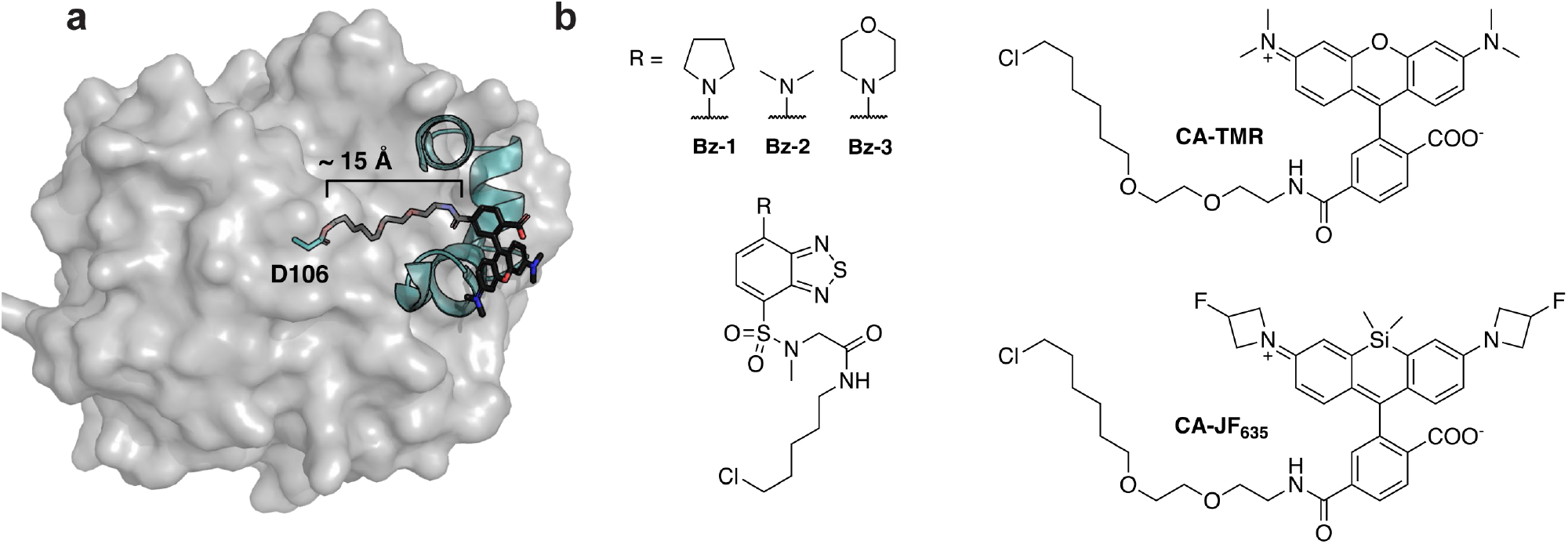
HaloTag7 and dye-containing substrates. (a) Crystal structure of HaloTag7 covalently reacted with substrate chloroalkane-tetramethylrhodamine (**CA-TMR**, PDB: 6Y7A).^16^ The model highlights how the catalytic residue, D106, is at the bottom of a ∼15Å hydrophobic channel. This model also highlights how interactions between the rhodamine dye and the surface helices of HaloTag7 drive binding, illustrating why non-rhodamine substrates often have slower kinetics. (b) Structures of fluorogenic dyes, **Bz-1, Bz-2** and **Bz-3**, and structures of other common HaloTag7 ligands, **CA-TMR** and **CA-JF**_**635**_.

HaloTag7 has proven to be very versatile given its genetic encodability and substrate modularity. However, its reaction rate with chloroalkane-tagged substrates is highly variable and substrate-specific.^16^ Specifically, HaloTag7 reacts with chloroalkane-tetramethylrhodamine (**CA-TMR**, Fig. 1) with second-order rate constants greater than 10^7^ M^-1^s^-1^ but this rate decreases substantially when non-rhodamine dyes are used. For example, the chloroalkane conjugate of AlexaFluor 488, a synthetic derivative of fluorescein that still bears a xanthene core, has a reaction rate 3 orders of magnitude slower than that of **CA-TMR** (2.5×10^4^ M^-1^s^-1^).^16^ When non-xanthene dyes are used, like benzothiadiazoles or stilbenes,^17-19^ this rate further plummets below 10^3^ M^-1^s^-1^. HaloTag7’s preference for **CA-TMR** can be explained by the fact that **CA-TMR** was the substrate in the original HaloTag7 engineering efforts – HaloTag is no exception to the maxim “you get what you screen for.”^20, 21^ In cellular assays, slow reaction kinetics results in sluggish or incomplete HaloTag7 labeling^22^ and/or the need for higher concentration of dye in cellular experiments, which leads to high background fluorescence.^11^

HaloTag7’s substrate bias has been accommodated by using rhodamines and other xanthene-based dyes that range in spectral properties.^23-27^ Fluorogenic HaloTag systems, which use dyes that become fluorescent upon conjugation with HaloTag7, are especially attractive as they increase sensitivity and eliminate the need for washout steps, allowing for real-time monitoring of biochemical processes.^28-30^ However, many of fluorogenic HaloTag7 ligands deviate from the rhodamine scaffold and thus suffer from slow reaction rates.^17-19, 31, 32^ Rhodamine-based fluorogenic dyes such as **CA-JF**_**635**_ (Fig. 1b) have also been developed, and their conjugation kinetics are faster than non-rhodamine dyes but they do not approach the super-fast kinetics of **CA-TMR**.^11, 16, 33, 34^ Given the broad utility of HaloTag7 in chemical biology, broadening its substrate scope to enable more rapid kinetics with non-xanthene dyes would permit a large expansion of the biosensor toolbox including the use of more varied fluorogenic dyes.

While most work to date has focused on improving dyes as substrates for HaloTag7, recent efforts to alter HaloTag7 to improve performance with a specific chloroalkane substrate have been reported. Liang, Ward, and coworkers screened a library of 73 recombinantly expressed and purified single-mutant HaloTag variants for improved activity of a catalytic metal center and, in a separate report, a similar library was screened for improved fluorogenic and labeling properties of a styrylpyridium dye.^35, 36^ Frei, Johnsson, and coworkers engineered HaloTag to modulate the fluorescent lifetimes of fluorogenic rhodamine dyes to enable multiplexed fluorescent lifetime imagining.^37^ They employed a HaloTag7 library generated by site-saturation mutagenesis of 10 pre-selected residues followed by screening bacterial lysates. Subsequent rounds of screening utilized sub-libraries generated by combinations of the best-performing single mutants. While both strategies produced improved HaloTag7 variants for their given application, they were limited by their screening throughput. In this work, we develop a molecular evolution system for HaloTag that can screen 10^7^ to 10^8^ variants for optimal properties including faster conjugation kinetics. The system was applied to produce an optimized HaloTag7 variant with improved kinetics for a fluorogenic benzothiadiazole dye. The new self-labeling protein, BenzoHTag, enables rapid wash-free intracellular labeling of live mammalian cells. Further, the new protein•dye system shows orthogonality to HaloTag7 which enabled both systems to be used simultaneously for multiplexed, wash-free labeling in live cells.

## Results

### Evolving HaloTag7 Using Yeast Surface Display

In contrast to previous methods for HaloTag7 evolution, we sought to employ a method that would enable higher throughput. We adapted yeast surface display for this purpose,^38-42^ which provides several benefits for protein evolution including the ability to sort using fluorescence-activated cell sorting and the inclusion of epitope tags to allow independent measurements of protein activity and expression level (Fig. 2a).^43^ HaloTag7 was incorporated into a yeast display construct and activity of HaloTag7 on the yeast surface was verified by treating yeast with **CA-TMR**. Robust **CA-TMR** signal was observed for yeast cells expressing HaloTag7 but not cells expressing the catalytically inactive D106A mutant (Fig. S1a). Immunostaining of HA and/or Myc epitopes produced a linear correlation with **CA-TMR** signal, demonstrating independent measurements of labeling activity and expression levels (Figs. 1a, S1b). This was important to avoid bias towards high-expressing variants in subsequent screens. We used error-prone PCR to generate four sub-libraries with 2 to 6 mutations per variant (Tables S3, S4). After verifying that each sub-library retained some activity (Fig. S2) they were pooled to yield an input library of 2.5×10^8^ variants. We filtered the input library to remove catalytically dead variants by treating a pool of over 4x10^9^ yeast with excess chloroalkane-biotin and then isolating biotinylated yeast using magnetic streptavidin beads. This pre-screen produced a filtered input library of functional HaloTag variants exceeding 5×10^7^ unique members.

**Figure 2.**
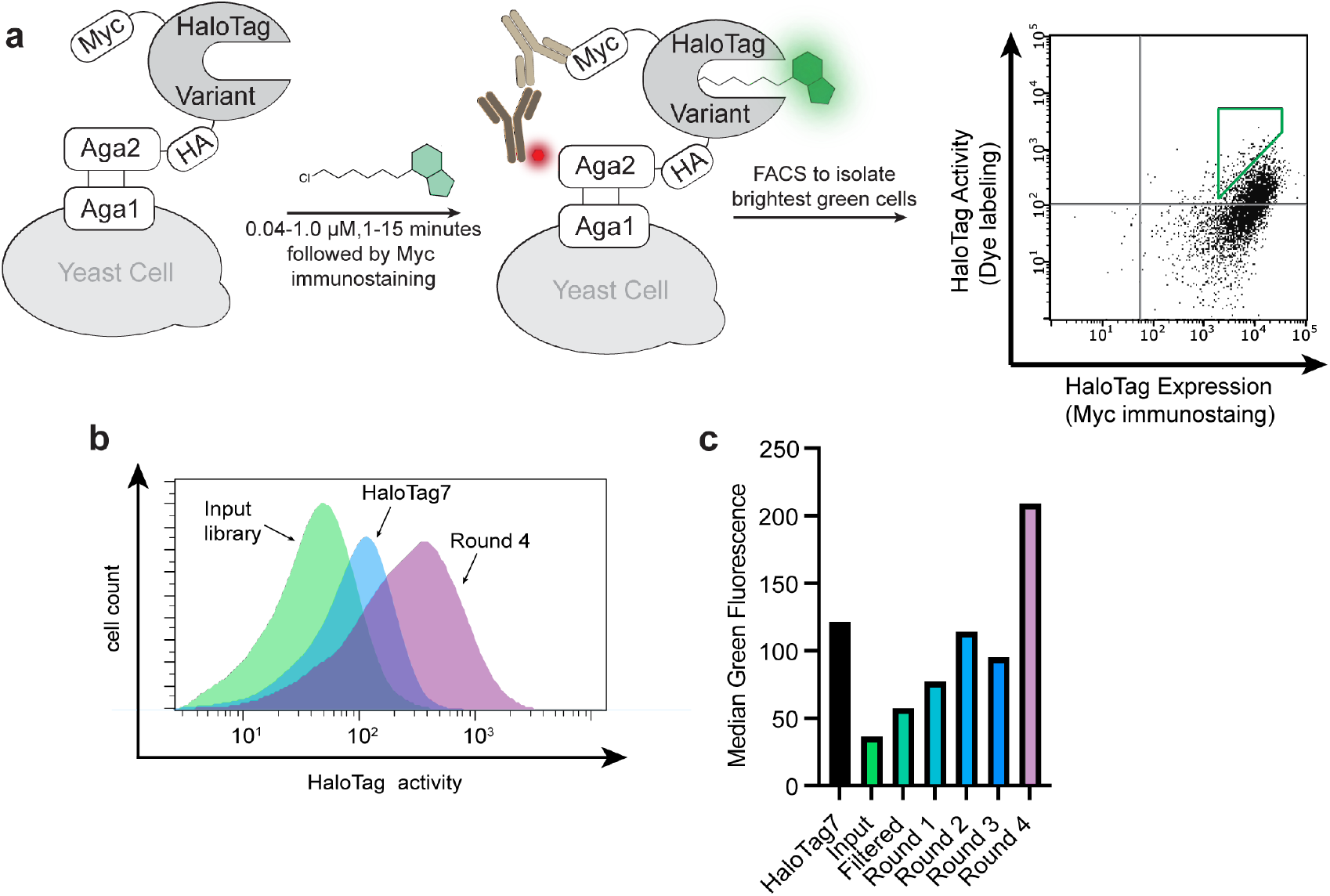
Molecular evolution of a fluorogenic HaloTag system using benzothiadiazole dye Bz-1. (a) Yeast display construct and screening strategy. Cells within the green gate were isolated and used for subsequent rounds of sorting. (b) Histograms of green fluorescence of 10,000 yeast cells displaying the input library (green), HaloTag7 (blue), or round 4 of the screen (purple). (c) Median green fluorescence of 10000 yeast cells displaying HaloTag7, the input library, the filtered input library, and the output pools of rounds 1 through rounds 4. Background green fluorescence from unlabeled cells was subtracted. Cells were incubated with 40 nM **Bz-1** for 1 minute.

To validate the HaloTag yeast display system, we screened the HaloTag variant library against **Bz-1**, a benzothiadiazole dye that we recently developed as a fluorogenic HaloTag ligand (Fig. 1b).^19^ Benzothiadiazoles are a class of fluorogenic dyes that are non-fluorescent in aqueous solution and fluorescent in non-polar environments, including the HaloTag7 active site channel as originally demonstrated by Zhang and coworkers.^17, 44^ **Bz-1** was cell-penetrant, it had low background in mammalian cells, and when conjugated to HaloTag7 **Bz-1** had spectral properties that align with GFP and AlexaFluor488 allowing for the use of common blue lasers and blue/green filter sets.^19, 45, 46^ Further, **Bz-1**’s small size, large Stokes shift of 70 nm which limits self-absorption, and ease of derivatization renders it a nearly ideal dye for *turn-on* fluorescence labeling in cells. Despite these favorable properties, the reaction rate of HaloTag7 conjugation to **Bz-1** was slower than the rates of many commonly used HaloTag7 substrates.^19^ Thus, we sought to improve the fluorogenic system by screening for HaloTag7 variants with improved reaction kinetics, and potentially improved fluorescence intensity with **Bz-1**. We subjected the filtered library to iterative rounds of screening using fluorescence-activated cell sorting with substrate **Bz-1**. In each round, we isolated the top 0.5% of cells with high green fluorescence relative to expression level (Fig. 2a). Stringency was increased after each round by decreasing the concentration of **Bz-1** and decreasing the incubation time, with round 4 applying 40 nM **Bz-1** for one minute (Table S6). After four rounds of screening, there was a clear increase in fluorescence of the sorted variants when treated with **Bz-1** compared to HaloTag7 (Fig. 2b-c).

### HaloTag mutations enhance conjugation with Bz-1

We sequenced 110 colonies from rounds 3 and 4. There were few duplicate sequences, but numerous enriched mutations were observed. 15 HaloTag variants that encompassed most of the enriched mutations were identified for further analysis. Notably, most enriched mutations were at residues that were not altered in prior HaloTag engineering efforts (Table S2)^5^ with the exception of V245A, which was identified in an earlier HaloTag7 evolution screen with a non-rhodamine, styrylpyridium dye.^36^ After comparing the activities of these 15 variants on the surface of yeast (Fig. S4 and S5, Table S7) we selected six of the best-performing variants for recombinant expression and purification (Variants 1-6, Table 1). All six variants demonstrated faster **Bz-1** labeling kinetics than HaloTag7 ((5-to 27-fold, Fig. 3a,b, Fig. S6, Table S10). The identified mutations also modulated the fluorescence properties of the protein•**Bz-1** complex (Fig. 3c). For example, Variant 1, which was the only variant to contain the L246F mutation, produced the largest enhancement in endpoint fluorescence intensity of 14% greater than HaloTag7. Further, all mutants bearing the V245A mutation produced a slightly red-shifted emission maximum (Table 1). Examination of a crystal structure of **Bz-2** conjugated with HaloTag7 suggested that all six variants have mutations that alter the environment near the dye’s benzothiadiazole core and/or donor amine group (Fig. 3d).^17, 36^ To explore whether these mutations altered interactions with **Bz-1**’s benzothiadiazole core or its donor amine, we compared the kinetics for Variants 5 and 6 reacting with **Bz-1, Bz-2**,^17^ and **Bz-3**,^19^ – these dyes have pyrrolidine, dimethylamine, and morpholine as their amine donors, respectively. The rates of **Bz-2** and **Bz-3** reacting with Variants 5 and 6 were approximately 10-fold slower than **Bz-1** but were approximately 10-fold faster than their rates with HaloTag7 (Fig. S7). These results implied that the newly evolved HaloTag variants specifically recognize both the benzothiadiazole core and the pyrrolidine donor group of **Bz-1**.

**Table 1.**
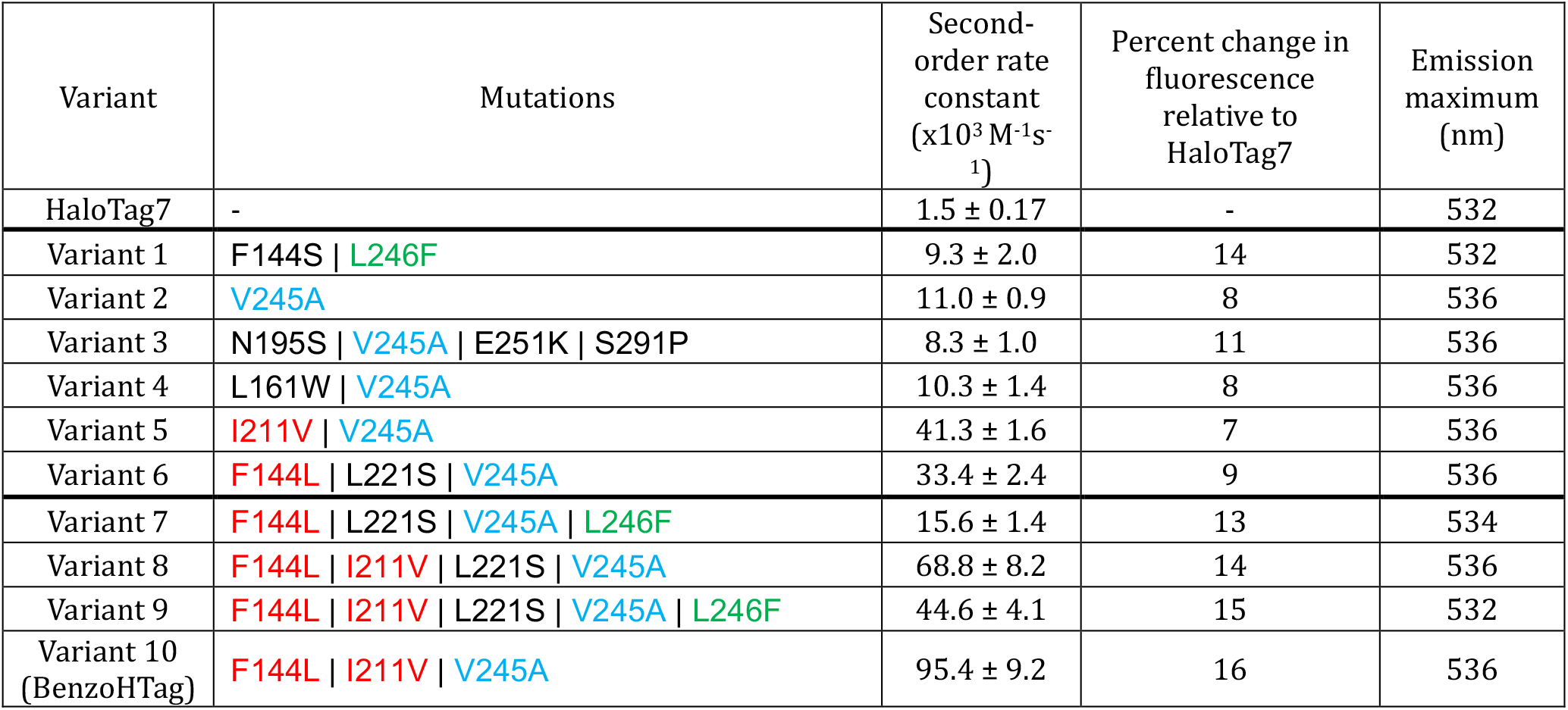
Summary of HaloTag Variant Sequences.

| Variant | Mutations | Second-order rate constant ($\times 10^3 \text{ M}^{-1} \text{ s}^{-1}$) | Percent change in fluorescence relative to HaloTag7 | Emission maximum (nm) |
| --- | --- | --- | --- | --- |
| HaloTag7 | - | $1.5 \pm 0.17$ | - | 532 |
| Variant 1 | F144S \| L246F | $9.3 \pm 2.0$ | 14 | 532 |
| Variant 2 | V245A | $11.0 \pm 0.9$ | 8 | 536 |
| Variant 3 | N195S \| V245A \| E251K \| S291P | $8.3 \pm 1.0$ | 11 | 536 |
| Variant 4 | L161W \| V245A | $10.3 \pm 1.4$ | 8 | 536 |
| Variant 5 | I211V \| V245A | $41.3 \pm 1.6$ | 7 | 536 |
| Variant 6 | F144L \| L221S \| V245A | $33.4 \pm 2.4$ | 9 | 536 |
| Variant 7 | F144L \| L221S \| V245A \| L246F | $15.6 \pm 1.4$ | 13 | 534 |
| Variant 8 | F144L \| I211V \| L221S \| V245A | $68.8 \pm 8.2$ | 14 | 536 |
| Variant 9 | F144L \| I211V \| L221S \| V245A \| L246F | $44.6 \pm 4.1$ | 15 | 532 |
| Variant 10 (BenzoHTag) | F144L \| I211V \| V245A | $95.4 \pm 9.2$ | 16 | 536 |

**Figure 3.**
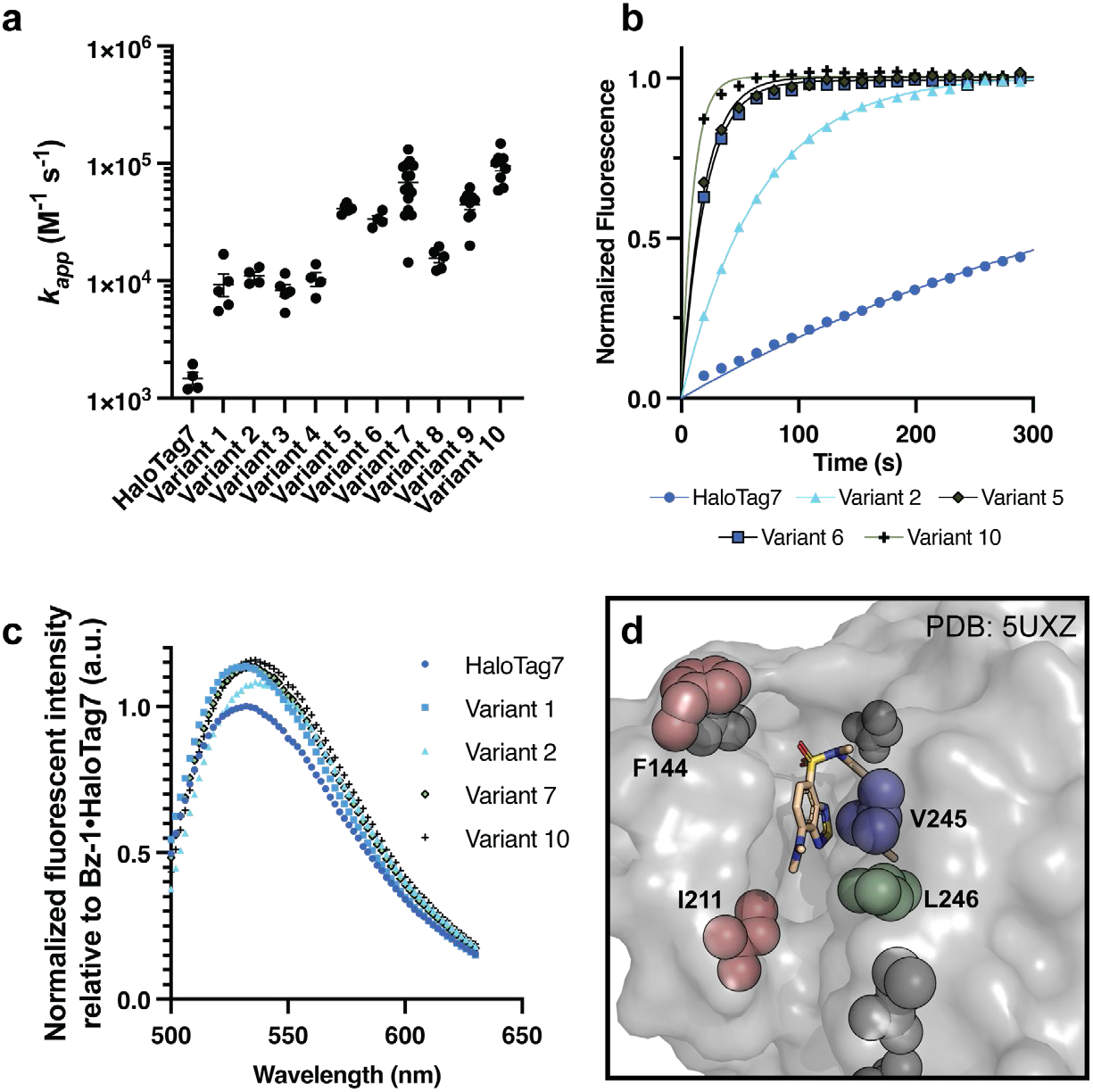
Characterization of recombinantly expressed HaloTag variants with improved fluorogenic properties. (a) Summary of second-order rate constants measured for **Bz-1** conjugation to HaloTag7 and Variants 1-10. See supplementary information for experimental details. (b) Representative kinetic traces of 0.25 μM **Bz-1** reacting with 1.0 μM HaloTag7 or selected variant. (c) Representative emission spectra of 2.5 μM **Bz-1** when conjugated to 5.0 μ? HaloTag7 or selected variant after one hour of incubation, normalized to the maximum fluorescence intensity of the HaloTag7•**Bz-1** complex. (d) Crystal structure of **Bz-2** with HaloTag7 (PDB: 5UXZ)^17^ showing the locations of key mutations observed in variants with improved fluorogenic properties.

We next generated Variants 7-10 that combined different mutations observed to correlate with rate enhancements among Variants 1-6. We observed that combining all three of the mutations V245A, F144L, and L211V (Variants 8-10) led to over two-fold faster rates compared to variants with only two of these mutations (Variants 5-7). We also observed that variants with the adjacent mutations V245A and L246F showed an overall decrease in reaction rate (Variants 7 and 9, compared to 6 and 8). Lastly L221S appeared to be a spectator mutation co-isolated with more beneficial mutations in Variant 6, as it is located distal to the active site channel and it decreases overall activity (Variant 10 compared to 8).

### Bz-1 rapidly labels BenzoHTag in live cells with low background

We selected Variant 10, which we named BenzoHTag, for testing in live mammalian cells. BenzoHTag was cloned as a Histone 2B (H2B) fusion to localize it to the nucleus and the fusion was transiently transfected into U-2 OS cells.^37^ Cells expressing BenzoHTag or HaloTag7 were treated for 10 minutes with concentrations of **Bz-1** between 7 and 1000 nM (Fig. 4a). Even at the highest concentration tested, **Bz-1** showed minimal background fluorescence in non-expressing cells. BenzoHTag dramatically outperformed HaloTag7 in live cell labeling with **Bz-1**, enabling robust fluorescence at 10-to 20-fold lower concentration of **Bz-1**. Saturation of the turn-on signal was not observed for HaloTag7-expressing cells even at 1000 nM **Bz-1**, while turn-on signal saturated for BenzoHTag at 250 nM. Between 30 and 250 nM, BenzoHTag-expressing cells had 5-to 8-fold higher fluorescence over background compared to HaloTag7-expressing cells. BenzoHTag•**Bz-1** labeling was also very sensitive – after labeling for 10 minutes with only 7 nM **Bz-1**, BenzoHTag-expressing cells showed greater than 200-fold signal over background (Fig. 4a). These results highlights that the intrinsic properties of the **Bz-1** dye, including high cell permeability of the **Bz-1** substrate and very low background fluorescence,^19^ synergize with the increased reaction rate to allow robust fluorescence detection at very low dye concentrations.

**Figure 4.**
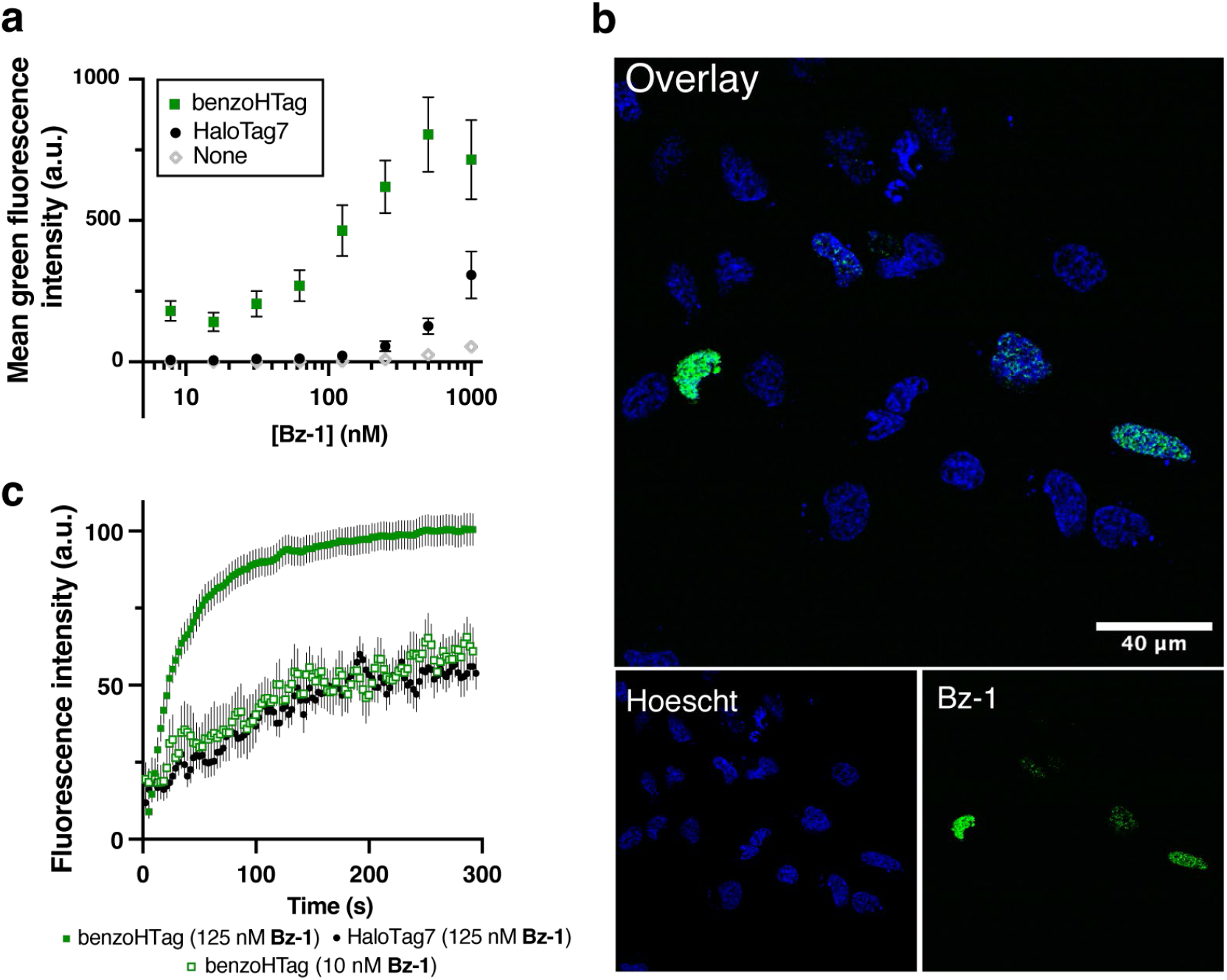
Comparing BenzoHTag and HaloTag7 in no-wash, live-cell labeling experiments. (a) Labeling with varied concentrations of **Bz-1** for 10 minutes in non-transfected U-2 OS cells and cells transfected with H2B-BenzoHTag or H2B-HaloTag7 (transfection efficiency was roughly 20%, Fig. S9). (b) Confocal microscopy imaging of H2B-BenzoHTag-transfected U-2 OS cells treated with 10 nM **Bz-1**. Cells were co-stained with Hoescht to highlight the nucleus. (c) Increase of **Bz-1** fluorescence over time when added to U-2 OS cells that were transiently transfected with H2B-BenzoHTag or H2B-HaloTag7. Fluorescence intensities of at least 10 transfected cells for each condition were measured over the course of five minutes with the average background fluorescence of 10 non-transfected cells subtracted. See Supporting Information for details.

We next evaluated the performance of the BenzoHTag•**Bz-1** system in no-wash, live cell fluorescence microscopy. U-2 OS cells transfected with the H2B-BenzoHTag fusion were treated with 10 nM **Bz-1** and imaged without exchanging media. Robust nuclear labeling was observed in BenzoHTag-expressing cells (Fig. 4b) while non-expressing cells within the same image had no observable background fluorescence. When treated with 125 nM **Bz-1**, cells also showed strong nuclear labeling with no detectable non-specific fluorescence (Fig. S11). We captured movies of **Bz-1**-treated cells (Movies S1, S2) and quantified the appearance of fluorescence over time. In BenzoHTag-expressing cells, fluorescence approached saturation within 60 seconds of **Bz-1** addition, and BenzoHTag-expressing cells saturated at 100% higher fluorescence intensities than HaloTag7-expressing cells (Fig. 4c, Fig. S12). Notably, BenzoHTag-expressing cells treated with only 10 nM **Bz-1** were also labeled within seconds and showed in-cell labeling kinetics similar to HaloTag7-expressing cells treated with 125 nM **Bz-1**. By contrast, no signal could be detected for HaloTag7-expressing cells when treated with only 10 nM **Bz-1**.

### BenzoHTag and HaloTag7 can be used for simultaneous, multiplexed labeling in live cells

Given that BenzoHTag recognizes multiple parts of **Bz-1**, we wondered whether the BenzoHTag system had evolved away from HaloTag7’s large preference for rhodamine-based substrates. To test this, we measured the kinetics of recombinantly purified BenzoHTag with **CA-TMR** and its fluorogenic silicon rhodamine analog, **CA-JF**_**635**_. **CA-TMR** reacted with BenzoHTag with a second-order rate constant of 2.1×10^4^ M^-1^s^-1^ and **CA-JF**_**635**_ reacted with a rate of 1.1×10^2^ M^-1^s^-1^, which represent 900- and 9000-fold rate decrease, respectively, relative to their rates with HaloTag7 (Fig. 5a, Fig. S8, Table S11). Overall, our kinetic data indicated that **Bz-1** reacts 65-fold faster with BenzoHTag than with HaloTag7, while **CA-JF**_**635**_ reacts 9000-fold faster with HaloTag7 than with BenzoHTag (See Supplemental Discussion). These results suggested that BenzoHTag•**Bz-1** and HaloTag7**•CA-JF**_**635**_ might be orthogonal enough for multiplexed labeling in cells. We then compared dye fluorescence in cells expressing either nucleus-localized BenzoHTag or HaloTag7. We observed that **CA-JF**_**635**_ preferentially labeled cells expressing HaloTag7 with very little labeling in cells expressing BenzoHTag, and **Bz-1** preferentially labeled cells expressing BenzoHTag with very little labeling in cells expressing HaloTag7 (Fig. 5b, S10)

**Figure 5.**
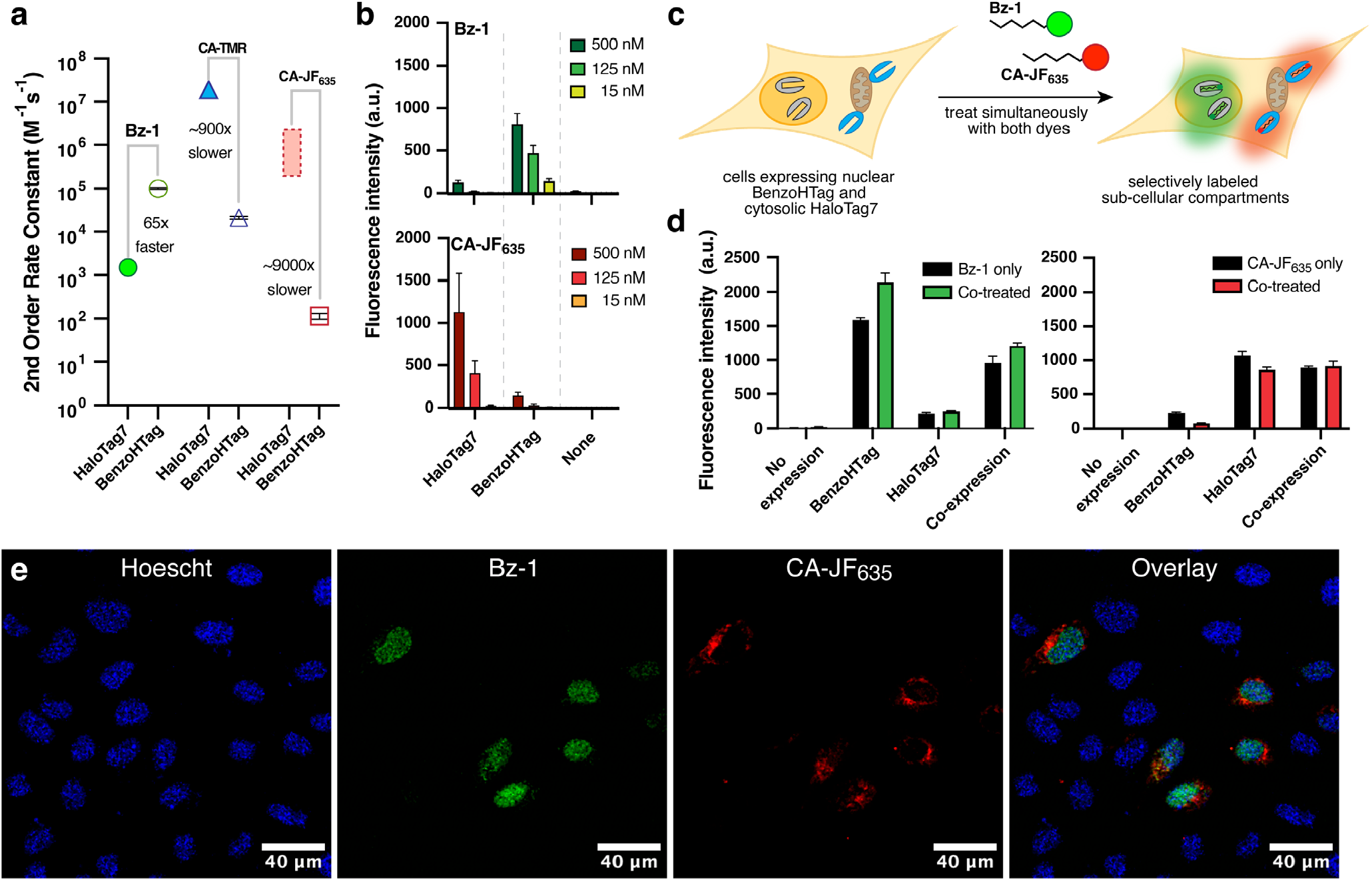
Multiplexed labeling using the BenzoHTag and HaloTag7 systems. (a) Second-order rate constants for **Bz1, CA-TMR**, and **CA-JF**_**635**_ with recombinantly expressed and purified HaloTag7 and BenzoHTag. The rate of **CA-TMR** with HaloTag7 was obtained and reported by Johnsson and coworkers.^16^ The rate of **CA-JF**_**635**_ has not been reported, but rates of analogous Si rhodamines were reported in the range of 10^5^-10^6^ M^-1^s^-1^.^11, 16, 23, 33^ See Supporting Information for more details. (b) Comparison of **Bz-1** and **CA-JF**_**635**_ labeling for 10 minutes in live U-2 OS cells transiently expressing H2B-BenzoHTag or H2B-HaloTag7. (c) Schematic of multiplexed labeling experiments. U-2 OS cells were transiently transfected with H2B-BenzoHTag (nuclear) and Tomm20-HaloTag7 (cytosolic, outer mitochondrial membrane fusion) and treated simultaneously with **Bz-1** and **CA-JF**_**635**_. (d) Flow cytometry data in orthogonal labeling experiments with 50 nM **Bz-1** and 125 nM **CA-JF**_**635**_ for 60 minutes. U-2 OS cells expressing either Tomm20-HaloTag7, H2B-BenzoHTag, or both were treated with either dye or co-treated with both dyes. The 15% most fluorescent cells were analyzed in each experiment because transient transfection efficiencies were roughly 20% (Fig. S9). (e) Confocal microscopy images of U-2 OS cells transiently transfected with both H2B-benzoHTag and Tomm20-HaloTag7. Cells were stained with nuclear Hoescht dye, washed, and then treated with 50 nM **Bz-1** and 125 nM **CA-JF**_**635**_ for 60 minutes. No washing was performed prior to imaging.

Encouraged by these results, we evaluated the ability to multiplex the BenzoHTag•**Bz-1** and HaloTag7•**CA-JF**_**635**_ systems in wash-free, live-cell labeling experiments. **Bz-1, CA-JF**_**635**_ or both dyes were added to U-2 OS cells transiently expressing BenzoHTag localized to the nucleus, HaloTag7 localized to the outer mitochondrial membrane, or both. Using 125 nM dye and analyzing cell populations by flow cytometry after 10 minutes of incubation, we observed that **Bz-1** predominately labeled BenzoHTag while **CA-JF**_**635**_ predominantly labeled HaloTag7 (Fig. S13b). To verify orthogonality under no-wash conditions in individual cells, we co-transfected cells with both constructs, treated them with both dyes simultaneously, and observed the cells using confocal fluorescence microscopy with no washes (Fig. 5c, Fig. S13c). When 125 nM of each dye was used, robust labeling by **Bz-1** was observed after 10 minutes but no **CA-JF**_**635**_ labeling was evident by microscopy. Imaging after 60 minutes revealed localization of **CA-JF**_**635**_ labeling to the mitochondria, however **Bz-1** labeling was observed at both the nucleus and mitochondria (Fig. S13c). We ascribe these observations to faster cell-penetration for **Bz-1** compared to **CA-JF**_**635**_. We optimized dye concentrations and found that co-treating cells with 50 nM **Bz-1** and 125 nM **CA-JF**_**635**_ for 60 minutes resulted in robust multiplexed labeling (Fig. 5d). **Bz-1** fluorescence was entirely localized to the nucleus and **CA-JF**_**635**_ fluorescence was entirely localized to the mitochondria, indicating excellent orthogonality between the BenzoHTag•**Bz-1** and HaloTag7•**CA-JF**_**635**_ systems (Fig. 5e). Quantification by flow cytometry under these conditions confirmed orthogonal labeling between the systems (Fig. 5d).

## Discussion

Self-labeling proteins like HaloTag7 have become a mainstay in chemical biology research. HaloTag7 is used for applications with a large variety of chloroalkane-tagged compounds, but the enzyme’s kinetics depend greatly on the nature of the substrate attached to the chloroalkane. Some prior work sought to modify HaloTag7 to improve the labeling rates with non-rhodamine substrates. An early example includes the mutation of negatively charged residues around the entrance to the active site channel of HaloTag to promote faster the conjugation of chloroalkane-tagged oligonucleotides.^47^ Most recently, several other groups have modified HaloTag7 to better accompany new substrates in a semi-rational approach using techniques that screen 10^2^-10^4^ variants at a time.^35-37, 48^ In this work, we developed a yeast display system capable of screening 10^7-108^ HaloTag7 variants at a time. We anticipate this system will greatly accelerate the development of HaloTag variants that work better with non-rhodamine substrates. Indeed, in this initial application, with a single round of diversification, the larger screening capability enabled the discovery of multiple cooperative mutations at unexpected positions; this result would have been highly unlikely using prior methods.^36, 37^ The yeast display format allows for a variety of positive and negative selections, rapid follow-up assays, and built-in controls for expression level.

In this first application, we provide ample evidence that yeast display produced HaloTag variants with improved conjugation kinetics and brighter fluorescent complexes. The optimized variant, BenzoHTag, had a 63-fold enhancement in reaction rate with **Bz-1** compared to HaloTag7. Our prior work optimized the fluorescent properties of the benzothiadiazole dye by replacing the dimethylamine donor in **Bz-2** with a pyrrolidine in **Bz-**1, which improved the fluorescent quantum yield of the dye when conjugated to HaloTag7 by 50% while decreasing the background in cells.^19^ Thus, combining dye engineering and protein engineering, we improved the original HaloTag7•**Bz-2** system by 1.5-fold in terms of brightness and by 300-fold in terms of reaction rate^17, 19^ We expect this roadmap – first optimizing the substrate for ideal functional properties, then optimizing the self-labeling protein for faster labeling – will enable additional systems to be developed with improved functionality compared to HaloTag7.

Prior to this work, the most well-developed fluorogenic substrate for HaloTag7 was a silicon rhodamine (SiR) dye, originally reported by Johnsson and colleagues^33^ and later optimized to CA-JF635 by Lavis and colleagues.^23, 34, 49^ We compared the performance of BenzoHTag•**Bz-1** and HaloTag7•**CA-JF**_**635**_ in live-cell labeling using both flow cytometry and fluorescence microscopy (Fig. 5). The reaction rates and relative brightness of the two systems were comparable in cells, but signal saturation was achieved much faster and at lower dye concentrations in the BenzoHTag•**Bz-1** system (Fig. 5b, S8). **CA-JF**_**635**_ is typically used in live-cell imaging experiments at concentrations of 100 nM to 500 nM, and sometimes exceeding 1 μM, with labeling times of 30 minutes or more.^23-25, 33, 34, 49^ These conditions match the concentrations and incubation times we observed were required for robust cellular labeling of HaloTag7•**CA-JF**_**635**._ By contrast, robust wash-free labeling was observed using only 10 nM **Bz-1** with an 18-fold signal-over-background as measured by confocal microscopy (Fig. S14), and nucleus-localized BenzoHTag was saturated by 125 nM **Bz-1** in under 100 seconds (Fig. 4c). When quantified by flow cytometry, **Bz-1** showed greater than 200-fold signal-over-background when applied at only 7 nM, (Fig. 4a, Fig. S8a-c). We interpret the differences in performance between HaloTag7•**CA-JF**_**635**_ and BenzoHTag•**Bz-1** to reflect superior cell permeability of the smaller, uncharged **Bz-1** compared to **CA-JF**_**635**_ (Fig. 1b). This interpretation is further supported by the observation that the percentage of cells labeled was not dependent on concentration of **Bz-1** but was highly dependent on concentration of **CA-JF**_**635**_ (Fig. S10c). The rapid labeling of the BenzoHTag•**Bz-1** system suggests unique for monitoring fast cellular processes, such as endosomal recycling.^11^ The no-wash nature of this system will also enable measurements of biological processes in real-time including protein folding in the cell, endocytosis and exocytosis rates, protein and organellar degradation processes, and endosomal escape of biological therapeutics.^13, 50-55^

There are several implementations of multiplexed fluorescent labeling using two different self-labeling proteins, like HaloTag7 and SNAP-tag,^23, 37, 56^ and even some reports of multiplexed no-wash fluorescent labeling.^57^ However, HaloTag7 is often preferred over SNAP-tag because SNAP-tag has slower reaction rates, its ligands have higher nonspecific interactions in the cell, and its complexes have weaker photophysical properties.^7, 16, 23, 58^ Therefore, it could be advantageous to use multiple HaloTag-derived self-labeling proteins in a wash-free multiplexed labeling experiments using orthogonal chloroalkane substrates. We found that BenzoHTag•**Bz-1** and HaloTag7•**CA-JF**_**635**_ support multiplexed no-wash labeling experiments in live cells (Fig. 5e). Given the ability to perform positive and negative selections using yeast display, we anticipate that additional HaloTag7 variants can be evolved for improved orthogonality and for specificity to other, spectrally orthogonal fluorogenic dyes that will allow multiplexing using three or more colors. Moreover, this strategy could be interfaced with recent advances in protein/peptide tags^59-61^ and fluorescence lifetime imaging^37, 62^ to offer even more degrees of multidimensional multiplexing.

## Conclusion

We have introduced the commonly used self-labeling protein HaloTag7 into a yeast display system for directed evolution of improved variants. This display platform can produce novel systems for imaging, biosensing, and biocatalysis that were previously inaccessible.^4, 5^ We used this system to develop BenzoHTag, an evolved HaloTag7 triple mutant that has improved conjugation kinetics to a fluorogenic benzothiadiazole dye, **Bz-1**. The BenzoHTag•**Bz-1** system enables robust intracellular labeling of live cells at concentrations as low as 7 nM, in seconds and without washes. The BenzoHTag•**Bz-1** system exhibits similar kinetics and maximum brightness as the previously reported HaloTag•**CA-JF**_**635**_ system but has faster and more sensitive in-cell labeling. The fast in-cell labeling rate will be especially useful for real-time monitoring of biological processes, especially intracellular processes, with fast time scales.^11, 54, 63^ Finally, the BenzoHTag•**Bz-1** system was found to be orthogonal to the HaloTag7•**CA-JF**_**635**_ system, allowing for simultaneous application of both systems for wash-free multiplexed imaging in live cells.

## Acknowledgments

This work was supported by NIH 5F32GM139371 to B.J.L. and NIH GM148407 to J.A.K. The authors would like to thank Prof. James Van Deventer for his thoughtful insights and for providing the RJY100 yeast strain, Dr. Luke Lavis for providing CA-JF_635_ samples, and Dr. Matt Lindley and the TAMIC core imaging facility for the confocal microscopy facilities, which is supported by the NIH infrastructure grant NIH S10 OD021624.

## TOC Graphic

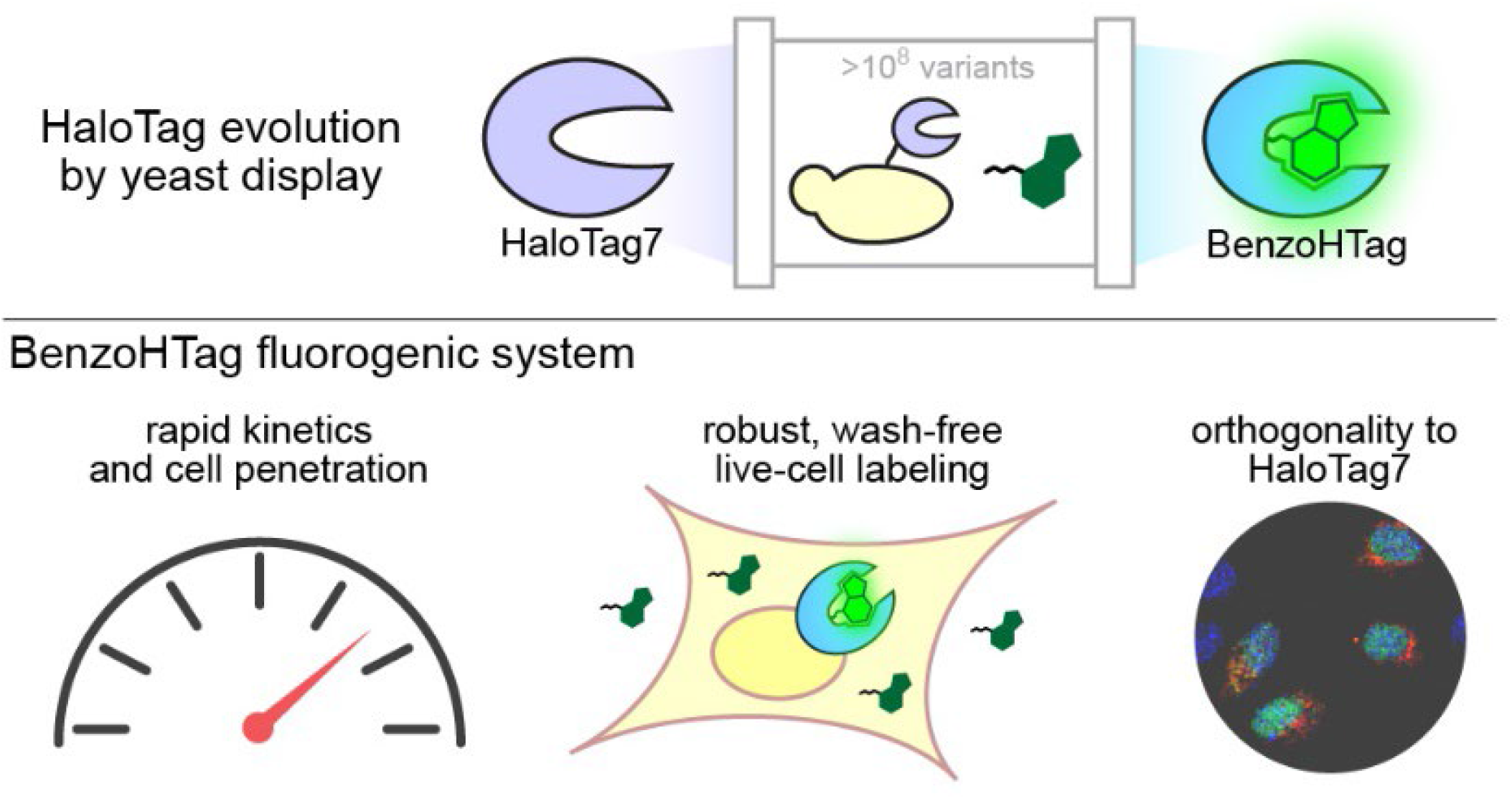

## References

(1) Greenwald, E. C.; Mehta, S.; Zhang, J. Genetically Encoded Fluorescent Biosensors Illuminate the Spatiotemporal Regulation of Signaling Networks. Chemical Reviews 2018, 118 (24), 11707–11794. DOI: 10.1021/acs.chemrev.8b00333.

(2) Wang, M.; Da, Y.; Tian, Y. Fluorescent proteins and genetically encoded biosensors. Chemical Society Reviews 2023, 52 (4), 1189–1214, 10.1039/D2CS00419D. DOI:10.1039/D2CS00419D.

(3) Los, G. V.; Encell, L. P.; McDougall, M. G.; Hartzell, D. D.; Karassina, N.; Zimprich, C.; Wood, M. G.; Learish, R.; Ohana, R. F.; Urh, M.; et al. HaloTag: A Novel Protein Labeling Technology for Cell Imaging and Protein Analysis. ACS Chemical Biology 2008, 3 (6), 373–382. DOI: 10.1021/cb800025k.

(4) England, C. G.; Luo, H.; Cai, W. HaloTag Technology: A Versatile Platform for Biomedical Applications. Bioconjugate Chemistry 2015, 26 (6), 975–986. DOI: 10.1021/acs.bioconjchem.5b00191.

(5) Cook, A.; Walterspiel, F.; Deo, C. HaloTag-Based Reporters for Fluorescence Imaging and Biosensing. ChemBioChem 2023, 24 (12), e202300022. DOI:10.1002/cbic.202300022 (acccessed 2023/09/10).

(6) Liss, V.; Barlag, B.; Nietschke, M.; Hensel, M. Self-labelling enzymes as universal tags for fluorescence microscopy, super-resolution microscopy and electron microscopy. Scientific Reports 2015, 5 (1), 17740. DOI:10.1038/srep17740.

(7) Erdmann, R. S.; Baguley, S. W.; Richens, J. H.; Wissner, R. F.; Xi, Z.; Allgeyer, E. S.; Zhong, S.; Thompson, A. D.; Lowe, N.; Butler, R.; et al. Labeling Strategies Matter for Super-Resolution Microscopy: A Comparison between HaloTags and SNAP-tags. Cell Chemical Biology 2019, 26 (4), 584-592.e586. DOI:10.1016/j.chembiol.2019.01.003.

(8) Deo, C.; Abdelfattah, A. S.; Bhargava, H. K.; Berro, A. J.; Falco, N.; Farrants, H.; Moeyaert, B.; Chupanova, M.; Lavis, L. D.; Schreiter, E. R. The HaloTag as a general scaffold for far-red tunable chemigenetic indicators. Nature Chemical Biology 2021, 17 (6), 718–723. DOI:10.1038/s41589-021-00775-w.

(9) Hellweg, L.; Edenhofer, A.; Barck, L.; Huppertz, M.-C.; Frei, M. S.; Tarnawski, M.; Bergner, A.; Koch, B.; Johnsson, K.; Hiblot, J. A general method for the development of multicolor biosensors with large dynamic ranges. Nature Chemical Biology 2023, 19 (9), 1147–1157. DOI:10.1038/s41589-023-01350-1.

(10) Shields, B. C.; Kahuno, E.; Kim, C.; Apostolides, P. F.; Brown, J.; Lindo, S.; Mensh, B. D.; Dudman, J. T.; Lavis, L. D.; Tadross, M. R. Deconstructing behavioral neuropharmacology with cellular specificity. Science 2017, 356 (6333), eaaj2161. DOI:10.1126/science.aaj2161 (acccessed 2023/09/10).

(11) Jonker, C. T. H.; Deo, C.; Zager, P. J.; Tkachuk, A. N.; Weinstein, A. M.; Rodriguez-Boulan, E.; Lavis, L. D.; Schreiner, R. Accurate measurement of fast endocytic recycling kinetics in real time. Journal of Cell Science 2020, 133 (2). DOI:10.1242/jcs.231225 (acccessed 3/29/2022).

(12) Machleidt, T.; Woodroofe, C. C.; Schwinn, M. K.; Méndez, J.; Robers, M. B.; Zimmerman, K.; Otto, P.; Daniels, D. L.; Kirkland, T. A.; Wood, K. V. NanoBRET—A Novel BRET Plaorm for the Analysis of Protein–Protein Interactions. ACS Chemical Biology 2015, 10 (8), 1797–1804. DOI:10.1021/acschembio.5b00143.

(13) Buckley, D. L.; Raina, K.; Darricarrere, N.; Hines, J.; Gustafson, J. L.; Smith, I. E.; Miah, A. H.; Harling, J. D.; Crews, C. M. HaloPROTACS: Use of Small Molecule PROTACs to Induce Degradation of HaloTag Fusion Proteins. ACS Chem Biol 2015, 10 (8), 1831–1837. DOI:10.1021/acschembio.5b00442From NLM.

(14) Peraro, L.; Deprey, K. L.; Moser, M. K.; Zou, Z.; Ball, H. L.; Levine, B.; Kritzer, J. A. Cell Penetration Profiling Using the Chloroalkane Penetration Assay. Journal of the American Chemical Society 2018, 140 (36), 11360–11369. DOI:10.1021/jacs.8b06144.

(15) Deprey, K.; Batistatou, N.; Debets, M. F.; Godfrey, J.; VanderWall, K. B.; Miles, R. R.; Shehaj, L.; Guo, J.; Andreucci, A.; Kandasamy, P.; et al. Quantitative Measurement of Cytosolic and Nuclear Penetration of Oligonucleotide Therapeutics. ACS Chemical Biology 2022, 17 (2), 348–360. DOI:10.1021/acschembio.1c00830.

(16) Wilhelm, J.; Kühn, S.; Tarnawski, M.; Gotthard, G.; Tünnermann, J.; Tänzer, T.; Karpenko, J.; Mertes, N.; Xue, L.; Uhrig, U.; et al. Kinetic and Structural Characterization of the Self-Labeling Protein Tags HaloTag7, SNAP-tag, and CLIP-tag. Biochemistry 2021. DOI:10.1021/acs.biochem.1c00258.

(17) Liu, Y.; Miao, K.; Dunham, N. P.; Liu, H.; Fares, M.; Boal, A. K.; Li, X.; Zhang, X. The Cation−π Interaction Enables a Halo-Tag Fluorogenic Probe for Fast No-Wash Live Cell Imaging and Gel-Free Protein Quantification. Biochemistry 2017, 56 (11), 1585–1595. DOI:10.1021/acs.biochem.7b00056.

(18) Clark, S. A.; Singh, V.; Vega Mendoza, D.; Margolin, W.; Kool, E. T. Light-Up “Channel Dyes” for Haloalkane-Based Protein Labeling in Vitro and in Bacterial Cells. Bioconjugate Chemistry 2016, 27 (12), 2839–2843. DOI:10.1021/acs.bioconjchem.6b00613.

(19) Lampkin, B. J.; Kritzer, J. A. Engineered fluorogenic HaloTag ligands for turn-on labelling in live cells. Chemical Communications 2024, 60 (2), 200–203. DOI:10.1039/D3CC05536A.

(20) Encell, L. P.; Friedman Ohana, R.; Zimmerman, K.; Otto, P.; Vidugiris, G.; Wood, M. G.; Los, G. V.; McDougall, M. G.; Zimprich, C.; Karassina, N.; et al. Development of a dehalogenase-based protein fusion tag capable of rapid, selective and covalent attachment to customizable ligands. Curr Chem Genomics 2012, 6, 55–71. DOI:10.2174/1875397301206010055From NLM.

(21) Marques, S. M.; Slanska, M.; Chmelova, K.; Chaloupkova, R.; Marek, M.; Clark, S.; Damborsky, J.; Kool, E. T.; Bednar, D.; Prokop, Z. Mechanism-Based Strategy for Optimizing HaloTag Protein Labeling. JACS Au 2022. DOI:10.1021/jacsau.2c00002.

(22) Thevathasan, J. V.; Kahnwald, M.; Cieśliński, K.; Hoess, P.; Peneti, S. K.; Reitberger, M.; Heid, D.; Kasuba, K. C.; Hoerner, S. J.; Li, Y.; et al. Nuclear pores as versatile reference standards for quantitative superresolution microscopy. Nature Methods 2019, 16 (10), 1045–1053. DOI:10.1038/s41592-019-0574-9.

(23) Grimm, J. B.; Tkachuk, A. N.; Xie, L.; Choi, H.; Mohar, B.; Falco, N.; Schaefer, K.; Patel, R.; Zheng, Q.; Liu, Z.; et al. A general method to optimize and functionalize red-shied rhodamine dyes. Nature Methods 2020, 17 (8), 815–821. DOI:10.1038/s41592-020-0909-6.

(24) Lardon, N.; Wang, L.; Tschanz, A.; Hoess, P.; Tran, M.; D’Este, E.; Ries, J.; Johnsson, K. Systematic Tuning of Rhodamine Spirocyclization for Super-resolution Microscopy. Journal of the American Chemical Society 2021, 143 (36), 14592–14600. DOI:10.1021/jacs.1c05004.

(25) Wang, L.; Tran, M.; D’Este, E.; Roberti, J.; Koch, B.; Xue, L.; Johnsson, K. A general strategy to develop cell permeable and fluorogenic probes for multicolour nanoscopy. Nature Chemistry 2020, 12 (2), 165–172. DOI:10.1038/s41557-019-0371-1.

(26) Zheng, Q.; Ayala, A. X.; Chung, I.; Weigel, A. V.; Ranjan, A.; Falco, N.; Grimm, J. B.; Tkachuk, A. N.; Wu, C.; Lippincott-Schwartz, J.; et al. Rational Design of Fluorogenic and Spontaneously Blinking Labels for Super-Resolution Imaging. ACS Central Science 2019, 5 (9), 1602–1613. DOI:10.1021/acscentsci.9b00676.

(27) Si, D.; Li, Q.; Bao, Y.; Zhang, J.; Wang, L. Fluorogenic and Cell-Permeable Rhodamine Dyes for High-Contrast Live-Cell Protein Labeling in Bioimaging and Biosensing. Angewandte Chemie International Edition 2023, 62 (45), e202307641. DOI:10.1002/anie.202307641.

(28) Prii, E.; Reymond, L.; Umebayashi, M.; Hovius, R.; Riezman, H.; Johnsson, K. A Fluorogenic Probe for SNAP-Tagged Plasma Membrane Proteins Based on the Solvatochromic Molecule Nile Red. ACS Chemical Biology 2014, 9 (3), 606–612. DOI:10.1021/cb400819c.

(29) Li, C.; Tebo, A. G.; Gautier, A. Fluorogenic Labeling Strategies for Biological Imaging. Int J Mol Sci 2017, 18 (7). DOI:10.3390/ijms18071473From NLM.

(30) Chen, Y.; Jiang, H.; Hao, T.; Zhang, N.; Li, M.; Wang, X.; Wang, X.; Wei, W.; Zhao, J. Fluorogenic Reactions in Chemical Biology: Seeing Chemistry in Cells. Chemical & Biomedical Imaging 2023. DOI:10.1021/cbmi.3c00029.

(31) Bachollet, S. P. J. T.; Shpinov, Y.; Broch, F.; Benaissa, H.; Gautier, A.; Pietrancosta, N.; Mallet, J.-M.; Dumat, B. An expanded palette of fluorogenic HaloTag probes with enhanced contrast for targeted cellular imaging. Organic & Biomolecular Chemistry 2022, 20 (17), 3619-3628, 10.1039/D1OB02394B. DOI:10.1039/D1OB02394B.

(32) Bachollet, S. P. J. T.; Addi, C.; Pietrancosta, N.; Mallet, J.-M.; Dumat, B. Fluorogenic Protein Probes with Red and Near-Infrared Emission for Genetically Targeted Imaging**. Chemistry – A European Journal 2020, 26 (63), 14467–14473. DOI:10.1002/chem.202002911.

(33) Lukinavičius, G.; Umezawa, K.; Olivier, N.; Honigmann, A.; Yang, G.; Plass, T.; Mueller, V.; Reymond, L.; Corrêa Jr, I. R.; Luo, Z.-G.; et al. A near-infrared fluorophore for live-cell super-resolution microscopy of cellular proteins. Nature Chemistry 2013, 5 (2), 132–139. DOI:10.1038/nchem.1546.

(34) Grimm, J. B.; Muthusamy, A. K.; Liang, Y.; Brown, T. A.; Lemon, W. C.; Patel, R.; Lu, R.; Macklin, J. J.; Keller, P. J.; Ji, N.; et al. A general method to fine-tune fluorophores for live-cell and in vivo imaging. Nat Methods 2017, 14 (10), 987–994. DOI:10.1038/nmeth.4403From NLM.

(35) Fischer, S.; Ward, T. R.; Liang, A. D. Engineering a Metathesis-Catalyzing Artificial Metalloenzyme Based on HaloTag. ACS Catalysis 2021, 11 (10), 6343–6347. DOI:10.1021/acscatal.1c01470.

(36) Miró-Vinyals, C.; Stein, A.; Fischer, S.; Ward, T. R.; Deliz Liang, A. HaloTag Engineering for Enhanced Fluorogenicity and Kinetics with a Styrylpyridium Dye. ChemBioChem 2021. DOI:10.1002/cbic.202100424.

(37) Frei, M. S.; Tarnawski, M.; Roberti, M. J.; Koch, B.; Hiblot, J.; Johnsson, K. Engineered HaloTag variants for fluorescence lifetime multiplexing. Nature Methods 2021. DOI:10.1038/s41592-021-01341-x.

(38) Colby, D. W.; Kellogg, B. A.; Graff, C. P.; Yeung, Y. A.; Swers, J. S.; Wittrup, K. D. Engineering antibody affinity by yeast surface display. Methods Enzymol 2004, 388, 348–358. DOI:10.1016/s0076-6879(04)88027-3From NLM.

(39) Chen, I.; Dorr, B. M.; Liu, D. R. A general strategy for the evolution of bond-forming enzymes using yeast display. Proceedings of the National Academy of Sciences 2011, 108 (28), 11399–11404. DOI:10.1073/pnas.1101046108.

(40) Van Deventer, J. A.; Wittrup, K. D. Yeast Surface Display for Antibody Isolation: Library Construction, Library Screening, and Affinity Maturation. Humana Press, 2014; pp 151–181.

(41) Branon, T. C.; Bosch, J. A.; Sanchez, A. D.; Udeshi, N. D.; Svinkina, T.; Carr, S. A.; Feldman, J. L.; Perrimon, N.; Ting, A. Y. Efficient proximity labeling in living cells and organisms with TurboID. Nat Biotechnol 2018, 36 (9), 880–887. DOI:10.1038/nbt.4201From NLM.

(42) Plamont, M.-A.; Billon-Denis, E.; Maurin, S.; Gauron, C.; Pimenta, F. M.; Specht, C. G.; Shi, J.; Quérard, J.; Pan, B.; Rossignol, J.; et al. Small fluorescence-activating and absorption-shifting tag for tunable protein imaging in vivo. Proceedings of the National Academy of Sciences 2016, 113 (3), 497–502. DOI:10.1073/pnas.1513094113.

(43) Chao, G.; Lau, W. L.; Hackel, B. J.; Sazinsky, S. L.; Lippow, S. M.; Wittrup, K. D. Isolating and engineering human antibodies using yeast surface display. Nat Protoc 2006, 1 (2), 755–768. DOI:10.1038/nprot.2006.94From NLM.

(44) Neto, B. A. D.; Correa, J. R.; Spencer, J. Fluorescent Benzothiadiazole Derivatives as Fluorescence Imaging Dyes: A Decade of New Generation Probes. Chemistry – A European Journal 2022, 28 (4), e202103262, 10.1002/chem.202103262. DOI:10.1002/chem.202103262 (acccessed 2022/04/25).

(45) Ruhlandt, D.; Andresen, M.; Jensen, N.; Gregor, I.; Jakobs, S.; Enderlein, J.; Chizhik, A. I. Absolute quantum yield measurements of fluorescent proteins using a plasmonic nanocavity. Commun Biol 2020, 3 (1), 627. DOI:10.1038/s42003-020-01316-2From NLM.

(46) Lavis, L. D.; Raines, R. T. Bright Ideas for Chemical Biology. ACS Chemical Biology 2008, 3 (3), 142–155. DOI:10.1021/cb700248m.

(47) Koßmann, K. J.; Ziegler, C.; Angelin, A.; Meyer, R.; Skoupi, M.; Rabe, K. S.; Niemeyer, C. M. A Rationally Designed Connector for Assembly of Protein-Functionalized DNA Nanostructures. ChemBioChem 2016, 17 (12), 1102–1106. DOI:10.1002/cbic.201600039.

(48) Kang, M.-G.; Lee, H.; Kim, B. H.; Dunbayev, Y.; Seo, J. K.; Lee, C.; Rhee, H.-W. Structure-guided synthesis of a protein-based fluorescent sensor for alkyl halides. Chemical Communications 2017, 53 (66), 9226-9229, 10.1039/C7CC03714G. DOI:10.1039/C7CC03714G.

(49) Grimm, J. B.; English, B. P.; Chen, J.; Slaughter, J. P.; Zhang, Z.; Revyakin, A.; Patel, R.; Macklin, J. J.; Normanno, D.; Singer, R. H.; et al. A general method to improve fluorophores for live-cell and single-molecule microscopy. Nature Methods 2015, 12 (3), 244–250. DOI:10.1038/nmeth.3256.

(50) Samelson, A. J.; Bolin, E.; Costello, S. M.; Sharma, A. K.; O’Brien, E. P.; Marqusee, S. Kinetic and structural comparison of a protein’s cotranslational folding and refolding pathways. Science Advances 2018, 4 (5), eaas9098. DOI: doi:10.1126/sciadv.aas9098.

(51) Luo, N.; Yan, A.; Yang, Z. Measuring Exocytosis Rate Using Corrected Fluorescence Recovery Aer Photoconversion. Traffic 2016, 17 (5), 554–564. DOI:10.1111/tra.12380From NLM.

(52) Liu, Y.; Zhang, D.; Qu, Y.; Tang, F.; Wang, H.; Ding, A.; Li, L. Advances in Small-Molecule Fluorescent pH Probes for Monitoring Mitophagy. Chemical & Biomedical Imaging 2024, 2 (2), 81–97. DOI:10.1021/cbmi.3c00070.

(53) Koga, H.; Martinez-Vicente, M.; Macian, F.; Verkhusha, V. V.; Cuervo, A. M. A photoconvertible fluorescent reporter to track chaperone-mediated autophagy. Nature Communications 2011, 2 (1), 386. DOI:10.1038/ncomms1393.

(54) Sahni, A.; Qian, Z.; Pei, D. Cell-Penetrating Peptides Escape the Endosome by Inducing Vesicle Budding and Collapse. ACS Chemical Biology 2020, 15 (9), 2485–2492. DOI:10.1021/acschembio.0c00478.

(55) Lönn, P.; Kacsinta, A. D.; Cui, X. S.; Hamil, A. S.; Kaulich, M.; Gogoi, K.; Dowdy, S. F. Enhancing Endosomal Escape for Intracellular Delivery of Macromolecular Biologic Therapeutics. Sci Rep 2016, 6, 32301. DOI:10.1038/srep32301From NLM.

(56) Bottanelli, F.; Kromann, E. B.; Allgeyer, E. S.; Erdmann, R. S.; Wood Baguley, S.; Sirinakis, G.; Schepartz, A.; Baddeley, D.; Toomre, D. K.; Rothman, J. E.; et al. Two-colour live-cell nanoscale imaging of intracellular targets. Nature Communications 2016, 7 (1), 10778. DOI:10.1038/ncomms10778.

(57) Martin, A.; Rivera-Fuentes, P. A general strategy to develop fluorogenic polymethine dyes for bioimaging. Nature Chemistry 2024, 16 (1), 28–35. DOI:10.1038/s41557-023-01367-y.

(58) Presman, D. M.; Ball, D. A.; Paakinaho, V.; Grimm, J. B.; Lavis, L. D.; Karpova, T. S.; Hager, G. L. Quantifying transcription factor binding dynamics at the single-molecule level in live cells. Methods 2017, 123, 76–88. DOI:10.1016/j.ymeth.2017.03.014From NLM.

(59) Gautier, A. Fluorescence-Activating and Absorption-Shifting Tags for Advanced Imaging and Biosensing. Accounts of Chemical Research 2022, 55 (21), 3125–3135. DOI:10.1021/acs.accounts.2c00098.

(60) Xue, L.; Karpenko, I. A.; Hiblot, J.; Johnsson, K. Imaging and manipulating proteins in live cells through covalent labeling. Nature Chemical Biology 2015, 11 (12), 917–923. DOI:10.1038/nchembio.1959.

(61) Suyama, A.; Devlin, K. L.; Macias-Contreras, M.; Doh, J. K.; Shinde, U.; Beatty, K. E. Orthogonal Versatile Interacting Peptide Tags for Imaging Cellular Proteins. Biochemistry 2023, 62 (11), 1735–1743. DOI:10.1021/acs.biochem.2c00712.

(62) Frei, M. S.; Koch, B.; Hiblot, J.; Johnsson, K. Live-Cell Fluorescence Lifetime Multiplexing Using Synthetic Fluorescent Probes. ACS Chemical Biology 2022, 17 (6), 1321–1327. DOI:10.1021/acschembio.2c00041.

(63) Komatsu, T.; Johnsson, K.; Okuno, H.; Bito, H.; Inoue, T.; Nagano, T.; Urano, Y. Real-Time Measurements of Protein Dynamics Using Fluorescence Activation-Coupled Protein Labeling Method. Journal of the American Chemical Society 2011, 133 (17), 6745–6751. DOI:10.1021/ja200225m.

