## Supplemental Information for "BenzoHTag, a fluorogenic self-labeling protein developed using molecular evolution"

### Supplementary Information

#### General Methods

All molecular biology reagents were purchased from New England Biolabs unless specified. Primers were ordered from Thermo Fisher Scientific. The HaloTag coding sequence was codon optimized for yeast expression and synthesized by Genscript. All recombinant cloning procedures were done by Gibson Assembly using 2x Master Mix from New England Biolabs following manufacturer's protocols. All antibodies used were purchased from Thermo Fisher Scientific. Yeast media supplements were purchased from Sigma Aldrich.

Kinetic and fluorescence endpoint assays were performed using a Tecan Spark plate reader. Flow cytometry experiments were conducted on a Guava cell analyzer and analyzed using Guava InCyte software. Fluorescence activated cell sorting (FACS) experiments were conducted on a BioRad S3E cell sorter. All microscopy was conducted on an Olympus IX71 fluorescence microscope and image acquisition and processing was done using CellSens Dimension software. Analysis of the of kinetics of live cell imaging was done using ImageJ and the Time Series Analyzer V3 plugin.[1]

#### Supplemental Discussion

*On BenzoHTag's decreased activity towards rhodamine substrates*

The three introduced mutations that yielded BenzoHTag decreased the conjugation kinetics of rhodamine-based **CA-TMR** and **CA-JF$_{635}$** by 900- and 9000-fold, respectively. HaloTag7 is the product of 25 mutations introduced into a haloalkane dehalogenase from *rhodococcus rhodochrous* (DhaA) that were introduced to improve both its labeling rate with the model substrate, **CA-TMR**, and improve its expression profile.[2, 3] Curiously, F144L, I211V, and V245A mutations are not at positions that were modified in the original HaloTag7 campaign (Table S2), posing the question why these mutations reduce activity towards rhodamine substrates. It has been computationally predicted that F144 stabilizes the *open* fluorescent isomer of a silicon rhodamine substrate,[4] so the F144L mutation may account for BenzoHTag's reduced activity towards rhodamine substrates. In fact, when measuring the kinetics of **CA-JF$_{635}$** conjugation to BenzoHTag, high concentrations (500 nM) of dye were needed to detect a *turn-on* in fluorescence upon reacting with BenzoHTag. This observation supports the critical role of F144 in rhodamine dye conjugation and promoting the isomerization to the *open* fluorescent isomer. Additionally, in work that screened site-saturation mutagenesis libraries of several positions, all mutants at F144 decreased fluorescence intensity with other fluorogenic rhodamine substrates that rely on the ground-state isomerization between a "closed" non-fluorescent spirolactone and an "open" fluorescent zwitterionic isomer.[5] This further supports a role for F144 in stabilizing the *open* fluorescent isomer for **CA-JF$_{635}$** and analogs conjugated to HaloTag7. A better understanding of the sequence-function relationship of HaloTag7 will better inform the relative reaction rates for HaloTag7 variants and more varied chloroalkane substrates.

**Table S1. Primer list**

| Name | |
|---|---|
| P1 | GGAGGCGGTAGCGGAGGCGGAGGGTCGGCTAGCTGCGGTGGCGGCGGTAT GGCTGAAATTGGTACAGGTTTTCCATTTG |
| P2 | GTCCTCTTCAGAAATAAGCTTTTGTTCGGATCCGCCCCCAGAAATTTCTAAAG |

| | |
|---|---|
| | TTGACAACCATCTAGCAATTTC |
| P3 | GTTCCAGACTACGCTCTGCAGG |
| P4 | CTAGTGGTGGAGGAGGCTCTGGTGGAGGCGGTAGCGGAGGCGGAGGGTCG GCTAGC |
| P5 | TATCAGATCTCGAGCTATTACAAGTCCTCTTCAGAAATAAGCTTTTGTTCGGA TCC |
| P6 | GCTCTGCAGGCTAGTGGTGGAGGAGGCTCTGGTG |
| P7 | CTACACTGTTGTTATCAGATCTCGAGCTATTACAAGTC |
| P8 | ATGGCTGAAATTGGTACAGGTTTTCCATTTG |
| P9 | AGAAATTTCTAAAGTTGACAACCATCTAGCAATTTC |
| P10 | AGAAGGAGATATACCATGCATCACCACCATCATCACATGGCTGAAATTGGTAC AGGTTTTC |
| P11 | AGCGGTGGCAGCAGCCTAGGTTAATTAAGAAATTTCTAAAGTTGACAACCATC |
| P12 | GGTAGCGGGGATCCACCGGTCGCCACCATGGCTGAAATTGGTACAGGTTTTC |
| P13 | AACGGGCCCTCTAGACTCGAGCGGTCAAGAAATTTCTAAAGTTGACAACCATC |
| P14 | ATGGCCAGAActgGCAAGAGAAA |
| P15 | TCATCCCATGTTGGAATTG |
| P16 | TGAATTACCAgtgGCTGGTGAAC |
| P17 | TTTGGAAATCTCCACAATG |
| P18 | ACCAGGTGCTTTTATTCCACCAG |
| P19 | CTGGTGGAATAAAAGCACCTGGT |

**Table S2. Summary of HaloTag mutations reported in previous work.**[a]

| | |
|---|---|
| Los*, Encel, Wood, et. al, *ACS Chem. Biol.,* **3**, 373 (2008) | |
| Purpose: convert dehalogenase to self-labeling enzyme | |
| HT | H272F |
| Encel*, Wood, et. al., Curr Chem Genomics., **6**, 55 (2012) | |
| Purpose: improve stability, expression, and kinetics with rhodamine dyes | |
| HT2 | K175M \| C176G \| H272F \| Y273L |
| HT3 | **HT2 +** S58T \| D78G \| A155T \| A172T \| A224E \| F272N \| P291S \| A292T \| plus Q294 \| plus Y295 |
| HT6 | **HT3 +** L47V \| Y87F \| L88M \| C128F \| E160K \| A167V \| K195N \| N227D \| E257K \| T264A |
| HT7 | **HT6 +** Q294E \| Y295I \| plus S296 \| plus G297 |
| Koßmann, Niemeyer*, et. al., ChemBioChem,17,1102 (2016) | |
| Purpose: improve HaloTag labeling with negatively charged substrates, like oligonucleotides | |
| HOB | **HT7 +** E154K \| E158K \| A166K \| D167K \| E171K |
| Liu, Zhang*, et. al., Angew. Chemie. Int. Ed., 56, 8672 (2017) | |
| Purpose: proteostasis sensor; destabllized HaloTag | |
| HOB | **HT7 +** K73T |
| Kang, Rhee*, et. al., Chem. Commun., 53, 9226 (2017) | |
| Purpose: Improve turn-on fluorescent properties of a dansyl dye | |
| $^{M175P}$HaloTag | **HT7 +** M175P |
| | ** also characterized M175T, N, K, C, H, Y, L, and W |

| | |
|---|---|
| Deprey & Kritzer*, *Bioconjugate Chemistry*, **32**, 964 (2021) | |
| Purpose: remove potential for inactivation via oxidation | |
| HT8 | **HT7 +** C61S \| C262S |
| Frei, Johnson*, et. al., *Nature Methods* **19**, 65 (2022) | |
| Purpose: modulate fluorescence lifetimes of rhodamine-based dyes | |
| HT9 | **HT7 +** Q165H \| P174R |
| HT10 | **HT7 +** Q165H |
| HT11 | **HT7 +** M175W |
| Miro-Vynyals, Liang*, et. al., *ChemBioChem*, **22**, 3398 (2021) | |
| Purpose: improve HaloTag labeling for a styrylpyridium dye | |
| HT-SP1 | **HT7 +** R133C \| M175Y \| V245A |
| HT-SP2 | **HT7 +** R133C \| E143M \| F144H \| M175Y \| V245A |
| HT-SP3 | **HT7 +** E143M \| F144A \| L271D |
| HT-SP4 | **HT7 +** M175Y \| L271D |
| HT-SP5 | **HT7 +** M175Y \| V245A \| L271D |
| This work | |
| Purpose: Improve brightness and kinetics with pro-fluorescent substrate | |
| Variant 1 | **HT7 +** F144S \| L246F |
| Variant 2 | **HT7 +** V245A |
| Variant 3 | **HT7 +** N195S \| V245A \| E251K \| S291P |
| Variant 4 | **HT7 +** L161W \| V245A |
| Variant 5 | **HT7 +** I211V \| V245A |
| Variant 6 | **HT7 +** F144L \| L221S \| V245A |
| Variant 7 | **HT7 +** F144L \| I211V \| L221S \| V245A |
| Variant 8 | **HT7 +** F144L \| L221S \| V245A \| L246F |
| Variant 9 | **HT7 +** F144L \| I211V \| L221S \| V245A \| L246F |
| Variant 10 **BenzoHTag** | **HT7 +** F144L + I211V + V245A |

[a]Further HaloTag engineering mutations, including split HaloTag and circularly permuted HaloTag, can be found in ref. 6

**Yeast Culture and Display Experiments**

Synthesized DNA encoding for HaloTag7 codon-optimized for *S. cerevisiae* was cloned into the linearized pCTcon2 (Gift from Dane Wittrup,[7] Addgene plasmid # 41843) construct between restriction sites NheI and BamHI using primers P1 and P2 to introduce homologous overhang regions.[7] Successfully cloned plasmid, determined by Sanger sequencing using primer P3, was used to transform the *S. cerevisiae* strain, RJY100 (Gift from James Van Deventer[8, 9]). Transformed yeast were selected using the TRP1 auxotrophic marker and were generally cultured according to previously published protocols.[10] Cells were propagated in synthetic dextrose media lacking tryptophan (SD-Trp, pH = 4.5) at 30 °C and grown to stationary phase. Approximately 24 hours prior to the display experiment, the $OD_{600}$ of the culture was measured, diluted to an $OD_{600}$ of 0.1 (~$1x10^7$ cells/mL), and grown to mid-log phase ($OD_{600}$ = 0.2-0.5, about 6 hours) at 30 °C. Cells were pelleted and then resuspended in synthetic galactose media with 2% dextrose and lacking tryptophan (SG-Trp, pH = 6.5) to an $OD_{600}$ of 0.1 to induce expression of Aga2-HaloTag fusion. Cells were induced at room temperature, shaking, for 16-18 hours prior to labeling.

For yeast display experiments, the induced culture's $OD_{600}$ was measured and $2x10^6$ cells were pelleted per sample. Cells were washed with three times with 50 μL with phosphate buffered saline (PBS) prior to labeling. To measure HaloTag activity on the surface of yeast, cells were resuspended in 50 μL of a solution of chloroalkane dye in PBS and incubated at room temperature for a set time. Cells were pelleted, dye solution was aspirated, then cells were washed three times with 50 μL with PBS. To measure HaloTag expression on the surface of yeast, primary and secondary antibody labeling of the epitope tags, HA and/or Myc, followed by a dye-labeled secondary antibody. Cells were resuspended in 50 μL of a 1:2000 diluted Rabbit anti-Myc and/or Mouse anti-HA in PBSA (PBS+0.1% w/v BSA) and tumbled at room temperature for 30 minutes. Cells were then pelleted at 4 °C and washed twice with 50 μL chilled PBSA. Cells were then resuspended in 50 μL of a 1:2000 diluted Goat anti-Rabbit labeled with AF647 and/or Goat anti-Mouse labeled with AF488 and kept on ice for 15 minutes. Cells were pelleted at 4 °C, washed once with 50 μL of chilled PBSA, resuspended in 200 μL of PBSA and transferred to a 96-well plate for flow cytometry analysis.

Yeast Codon-Optimized HaloTag genomic gene

ATGGCTGAAATTGGTACAGGTTTTCCATTTGATCCACATTACGTTGAAGTTTT
GGGTGAAAGAATGCATTACGTTGATGTTGGTCCAAGAGATGGTACACCAGT
TTTGTTTTTACATGGTAACCCAACTTCTTCATACGTTTGGAGAAACATCATCC
CACATGTTGCACCAACTCATAGATGTATTGCTCCAGATTTGATTGGTATGGG
TAAATCTGATAAGCCAGATTTGGGTTATTTCTTTGATGATCATGTTAGATTCA
TGGATGCTTTTATTGAAGCATTGGGTTTGGAAGAAGTTGTTTTGGTTATTCAT
GATTGGGGTTCTGCATTAGGTTTTCATTGGGCTAAGAGAAACCCAGAAAGA
GTTAAGGGTATCGCTTTTATGGAGTTTATTAGACCAATTCCAACATGGGATG
AATGGCCAGAATTTGCAAGAGAAACTTTCCAAGCTTTTAGAACAACTGATGT
TGGTAGAAAGTTGATCATCGATCAAAACGTTTTTATTGAAGGTACATTGCCA
ATGGGTGTTGTTAGACCATTGACTGAAGTTGAAATGGATCATTACAGAGAAC
CATTTTTAAACCCAGTTGATAGAGAACCATTGTGGAGATTTCCAAATGAATTA
CCAATTGCTGGTGAACCAGCAAACATCGTTGCTTTGGTTGAAGAATACATGG
ATTGGTTACATCAATCTCCAGTTCCAAAGTTGTTATTTTGGGGTACACCAGG
TGTTTTAATTCCACCAGCAGAAGCTGCAAGATTGGCTAAGTCATTGCCAAAC
TGTAAGGCAGTTGATATTGGTCCAGGTTTGAATTTGTTGCAAGAAGATAACC
CAGATTTGATTGGTTCTGAAATTGCTAGATGGTTGTCAACTTTAGAAATTTCT

Yeast Codon-Optimized HaloTag protein sequence

MAEIGTGFPFDPHYVEVLGERMHYVDVGPRDGTPVLFLHGNPTSSYVWRNIIP
HVAPTHRCIAPDLIGMGKSDKPDLGYFFDDHVRFMDAFIEALGLEEVVLVIHDW
GSALGFHWAKRNPERVKGIAFMEFIRPIPTWDEWPEFARETFQAFRTTDVGRK
LIIDQNVFIEGTLPMGVVRPLTEVEMDHYREPFLNPVDREPLWRFPNELPIAGEP
ANIVALVEEYMDWLHQSPVPKLLFWGTPGVLIPPAEAARLAKSLPNCKAVDIGP
GLNLLQEDNPDLIGSEIARWLSTLEIS

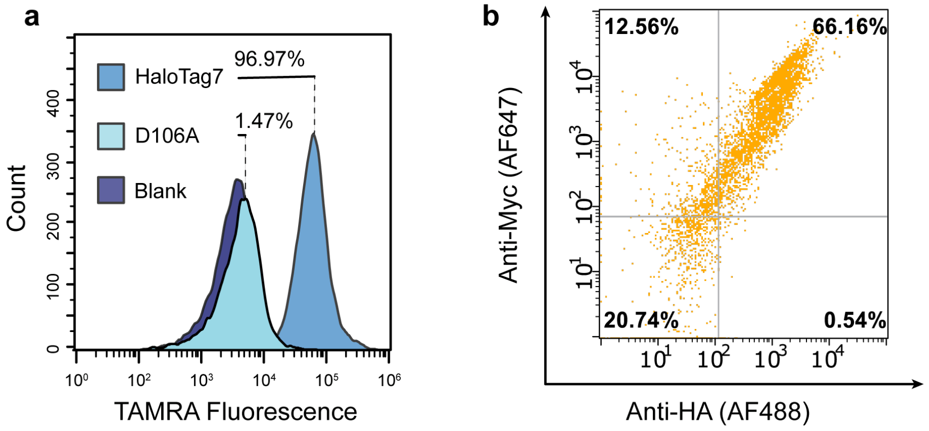

**Figure S1. HaloTag expression and activity on the surface of yeast**. a) Yeast that were induced to express HaloTag fusions at the surface were labeled with 1.0 µM chloroalkane-tetramethylrhodamine (TAMRA) for 30 minutes. Gating for uniform expression, HaloTag has near quantitative labeling while the catalytically inactive enzyme (97% of HaloTag7 cells were above background), D106A, displays no labeling. Blank cells refer to unlabeled cells. B) Immunostaining of the N and C terminal epitope tags, HA and Myc, respectively, indicates robust HaloTag expression on the surface of yeast after galactose induction.

**Combinatorial Library Generation**

Combinatorial HaloTag libraries were generated using error-prone PCR with mutagenetic nucleotides as described previously.[10-12] Briefly, HaloTag was amplified in a 50 µL reaction of 50 ng of template vector for 10-20 rounds using 0.2 µM of primers **P4** and **P5**, 2.5 mM $MgSO_4$, 5 units of *Taq* polymerase, 0.2 mM dNTPs, 2% DMSO, and 2 or 10 µM each of mutagenic nucleotide analogs 8-oxo-2'-deoxyguanosine-5'-triphosphate (8-oxo-dGTP) and 2'-deoxy-P-nucleoside-5'-triphosphate (dPTP) (Jena Biosciences, see Table S3). The PCR products were gel purified and amplified again under the same PCR conditions but without 8-oxo-dGTP and dPTP for 35 rounds using primers **P6** and **P7**.

**Table S3. Error-prone PCR Conditions**

| Entry | [8-oxo-dGTP], µM | [dPTP], µM | # of PCR cycles |
|---|---|---|---|
| 1 | 2 | 2 | 10 |
| 2 | 2 | 2 | 20 |
| 3 | 10 | 10 | 10 |
| 4 | 10 | 10 | 20 |

The inserts were electroporated into freshly made electrocompetent RJY100 with NheI-BamHI digested pCTcon2 vector in a 10:1 mass ratio following established protocols.[10] The electroporated cultures were rescued in 2 mL of yeast extract peptide dextrose media (YPD) for 1 hour at 30 °C without shaking. 10 µL of the recovered culture was used to generate 100x, 1000x, 10000x, and 100000x diluted samples of which 20 µL was plated on SD-Trp plates and incubated at 30 °C. Three days later, colony forming units were counted to estimate library size where one colony on 100x, 1000x, 10000x, and 100000x diluted plates corresponds to $1x10^4$, $1x10^5$, $1x10^6$, and $1x10^7$ transformants, respectively. Results are summarized in Table S4. The remaining recovered transformed culture was diluted in 100 mL of SD-Trp media and grown to saturation. This saturated culture was used to freeze library stocks that contained a number of cells corresponding to 10x the library diversity.

**Table S4. HaloTag Variant Libraries.**

| | Number of CFUs counted on diluted plates (# of transformants) | | | | |
|---|---|---|---|---|---|
| Sublibrary | 100x[a] | 1000x[a] | 10000x | 100000x | Average |
| 1 | N.D. | N.D. | 64 ($6.4x10^7$) | 8 ($8x10^7$) | $7.2x10^7$ |
| 2 | N.D. | N.D. | 116 ($1.16x10^8$) | 15 ($1.5x10^8$) | $1.33x10^8$ |
| 3 | N.D. | N.D. | 24 ($2.4x10^7$) | 2 ($2x10^7$) | $2.2x10^7$ |
| 4 | N.D. | N.D. | 19 ($1.9x10^7$) | 4 ($4x10^7$) | $2.95x10^7$ |

[a]*N.D* = Not determined; grown colonies were too dense to accurately count.

To evaluate amino acid mutation frequency, plasmid from each sublibrary was harvested by ZymoPrep (ZymoResearch), purified by precipitation, and re-transformed in NEB 10β *e. coli* cells by heat shock. Up to 24 random colonies from each sublibrary were inoculated into Luria-Bertani (LB) broth supplemented with 100 μg/mL ampicillin (GoldBio), plasmid was harvested by mini-prep (Invitrogen), and the insert region was by Sanger sequencing using **P3** to identify HaloTag variant sequences and mutation frequencies (Table S5).

**Table S5. Amino Acid Mutation Frequency in HaloTag Variant Sublibraries.**

| SubLibrary | Colonies sequenced | Average # of Mutations | Native HaloTag sequences detected |
|---|---|---|---|
| 1 | 19 | 2.9 | 3 |
| 2 | 21 | 2.2 | 2 |
| 3 | 24 | 6.4 | 0 |
| 4 | 21 | 9.5 | 0 |

The activity of each sublibrary was determined by flow cytometry (Fig. S2). Because each sublibrary appeared to have many active mutants, we pooled the four libraries together at 10x the sublibrary diversity to yield the input HaloTag variant library that has a diversity of $\sim 2.6x10^8$ total HaloTag variants. We estimated that <10% of this input library contains the native HaloTag sequence.

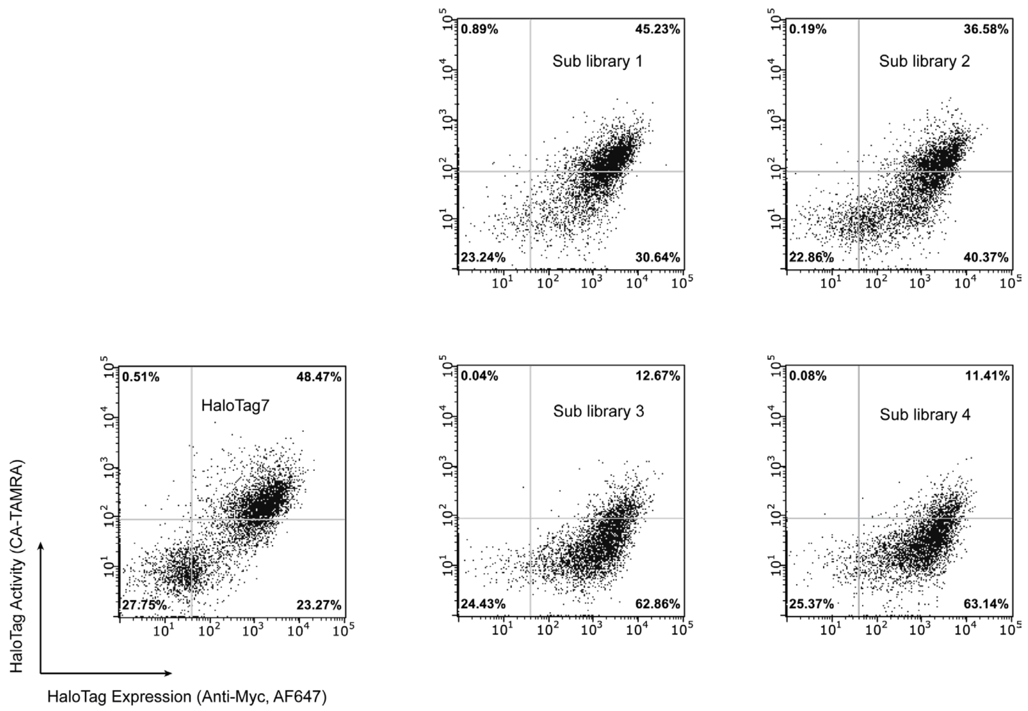

**Figure S2. HaloTag sublibrary activity**. Sublibraries were treated with 1.0 μM chloroalkane tetramethylrhodamine for 30 minutes.

**Yeast Cell Sorting**

*Magnetic* – The magnetic sort of the input library was done following a previously reported protocol.[10] Briefly, $4x10^9$ induced cells were pelleted (in four tubes of $1x10^9$ cells each) and each resuspended in 1 mL of PBSA. 20 μL of washed, streptavidin-coated magnetic beads were added to each tube of induced cells and tubes were tumbled at 4 °C for two hours. The tubes were placed a magnetic block, and the suspended cells were transferred to a fresh 2.0 mL tube and pelleted, thus removing cells that stuck nonspecifically to the magnetic beads. Cells were resuspended in 1.0 mL of 10 μM chloroalkane-biotin in PBS (estimated to be >10x excess of HaloTag copies) and tumbled at room temperature for one hour. The cells were then pelleted and washed three times with 1.0 mL PBS to remove excess ligand and resuspended in 1.0 mL of PBSA. Fresh, washed streptavidin-coated magnetic beads were added to the resuspended cells, tubes were tumbled at 4 °C for one hour, and then the tubes were place on a magnetic rack. The supernatant was carefully removed and the immobilized beads were gently washed twice with chilled PBSA, then pooled and suspended in 2.0 mL of SD-Trp. 10 μL was removed and used to make serial dilutions of beads (1:100, 1:1000, 1:10000, 1:100000) and 20 μL of each dilution was plated on a SD-Trp agar plate and incubated at 30 °C for 3 days prior to counting colonies. The remaining suspended beads were further diluted to 200 mL of SD-Trp media and grown to saturation. Based on colony counts, we estimated that this magnetic sort retained ~20% of the input cells yielding a filtered library of approximately $5.3x10^7$ active HaloTag variants.

*FACS* –1x10$^7$ yeast cell from the filtered library (or, later, from the previous round's output pool) were grown and induced as described above. All volumes for labeling and wash steps were increased to 250 µL to accommodate the 5x increase in labeled cells per tube. The labeling conditions of dye were varied for each round of FACS as described in Table S6. For FACS, labeled batches were pooled and resuspended to a final concentration of 1x10$^7$ cells/mL and subjected to sorting in which the top ~0.5% brightest green cells were isolated. Screening gates were shaped to include the brightest cells that were those well above the diagonal when green fluorescence (from the dye) was plotted against red fluorescence (from Myc immunostaining). Recovered cells were grown in SD-Trp media at 30 °C until saturation (~3 days) and the resulting stocks were then prepared for a subsequent round of sorting and frozen as glycerol stocks for later analysis.

**Table S6.** FACS labeling conditions

| | [**Bz-1**], µM | Time (min.) |
|---|---|---|
| Round 1 | 1.0 | 15 |
| Round 2 | 1.0 | 1 |
| Round 3 | 0.2 | 1 |
| Round 4 | 0.04 | 1 |

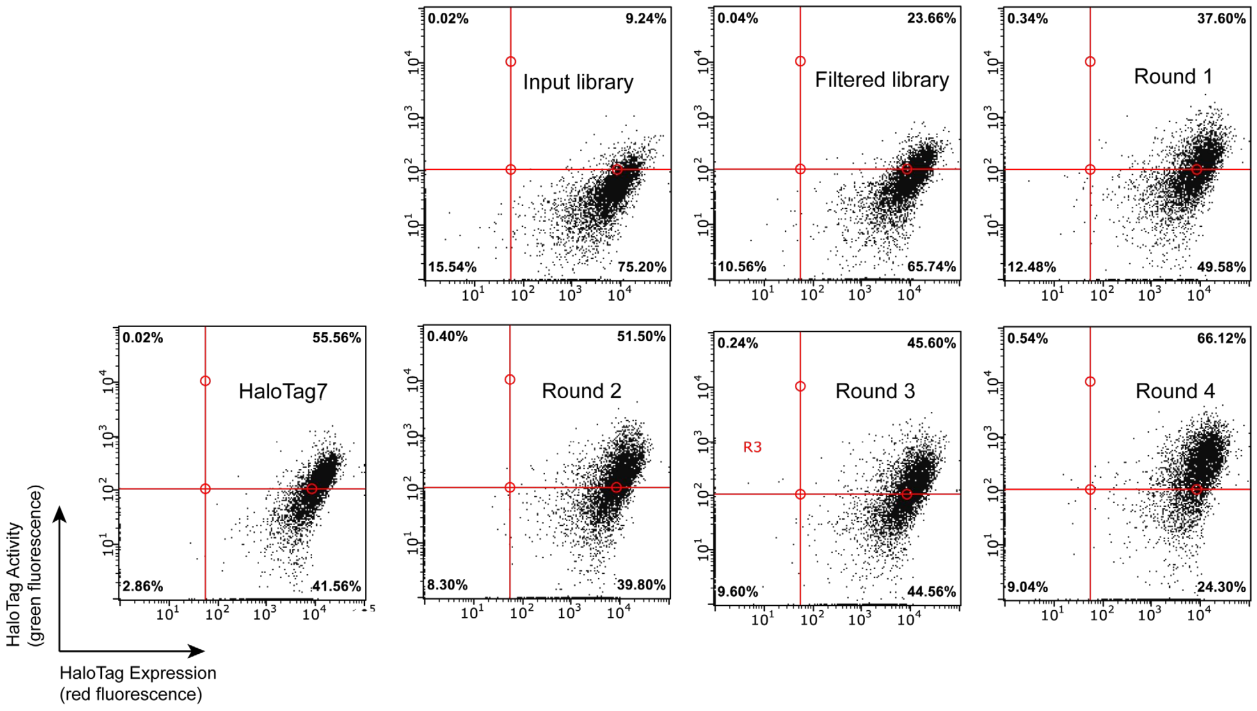

**Figure S3. Evaluating activity of the output pools from each round of screening.** HaloTag activity was measured by labeling with 1.0 µM **Bz-1** for 15 minutes. 10,000 cells are reported in each plot. HaloTag expression is measured by immunostaining against Myc using an AlexaFluor647-conjugated antibody.

After output pools round 3 and round 4 were isolated, 1x$10^8$ cells were pelleted and plasmid was harvested by ZymoPrep (ZymoResearch). Plasmids were purified, transformed, and sequenced as described above. Twenty colonies from R3 and 75 colonies from R4 were sequenced in total. Selected hit sequences were transformed into chemically competent RJY100 *S. cerevisiae* cells as described above, and HaloTag activity was evaluated by flow cytometry (Table S7, Fig. S4, S5). Six sequences were chosen for their high activity and/or representative mutations and their HaloTag variant sequences were isolated by PCR (20 ng of template, 0.2 μM primers **P8** and **P9,** 2.0 mM $MgSO_4$, 0.2 mM of each dNTP, 1 unit of Phusion High-Fidelity polymerase, and 2% DMSO) for cloning into bacterial expression vectors.

**Table S7**. Hit HaloTag Variants with Identified Mutations

| Variant | Alternate Name | Mutations |
|---|---|---|
| BzR3-1 | Variant 1 | F144S \| L246F |
| BzR3-5 | | D156N \| E191K \| V245A |
| BzR3-7 | | H13R |
| BzR3-12 | | E224K \| V245A |
| BzR3-20 | Variant 3 | N195S \| V245A \| E251K \| S291P |
| BzR3-21 | | L246F \| L259W |
| BzR4-11 | | F144L \| I211V \| V245A \| L271V |
| BzR4-27 | Variant 4 | L161W \| V245A |
| BzR4-42 | Variant 6 | F144L \| L221S \| V245A |
| BzR4-43 | Variant 5 | I211V \| V245A |
| BzR4-44 | Variant 2 | V245A |
| BzR4-45 | | D78G \| E98G \| E121G, V245A |
| BzR4-53 | | V245A \| A250V |
| BzR4-65 | | Q231R \| V245A |
| BzR4-67 | | A212V \| L246F |

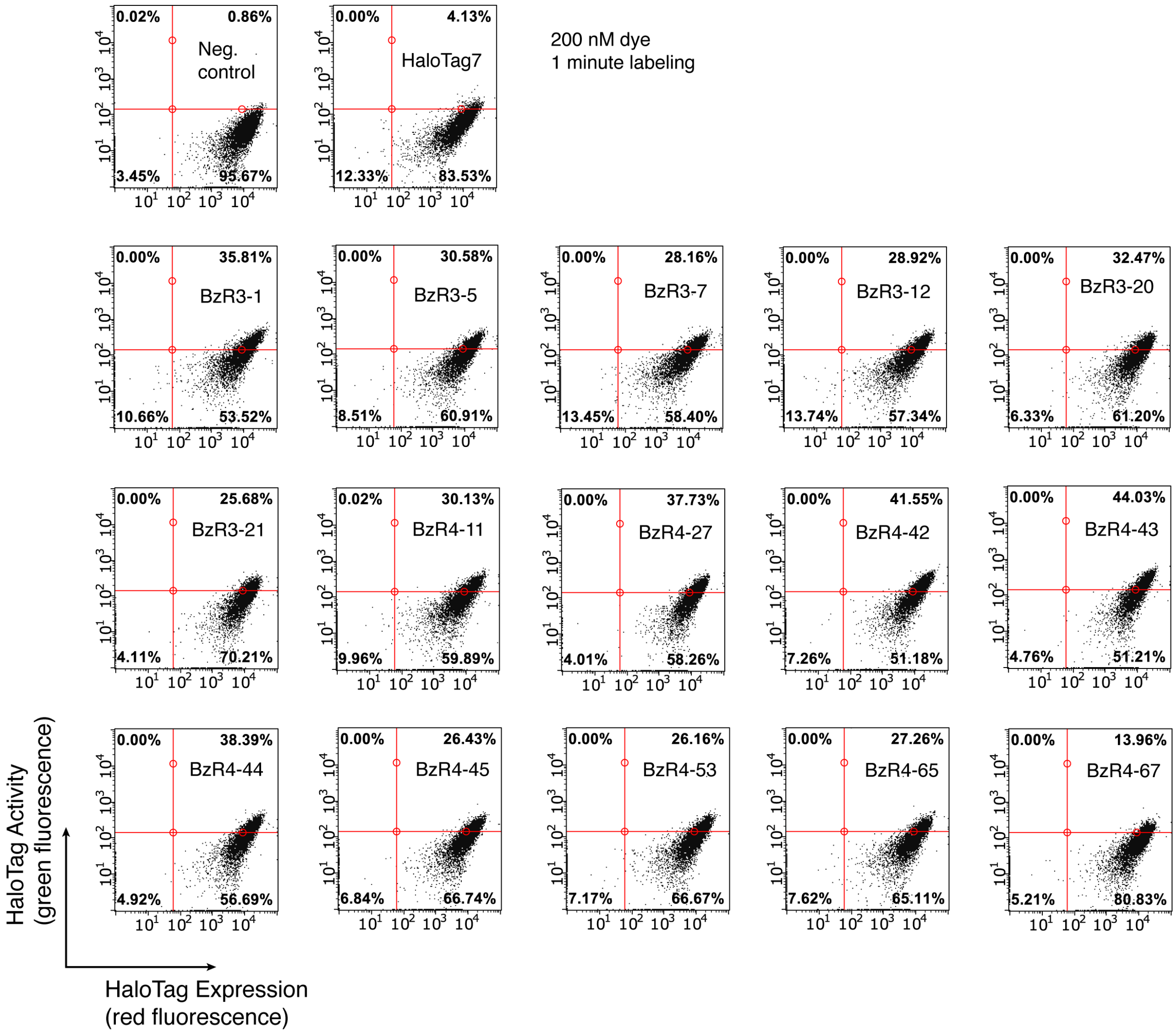

**Figure S4**. **Activity of screening hits, expressed on the surface of yeast analysis.** Single HaloTag variant clones were analyzed as described above. Cells were treated with 0.2 μM **Bz-1** for 1 minute.

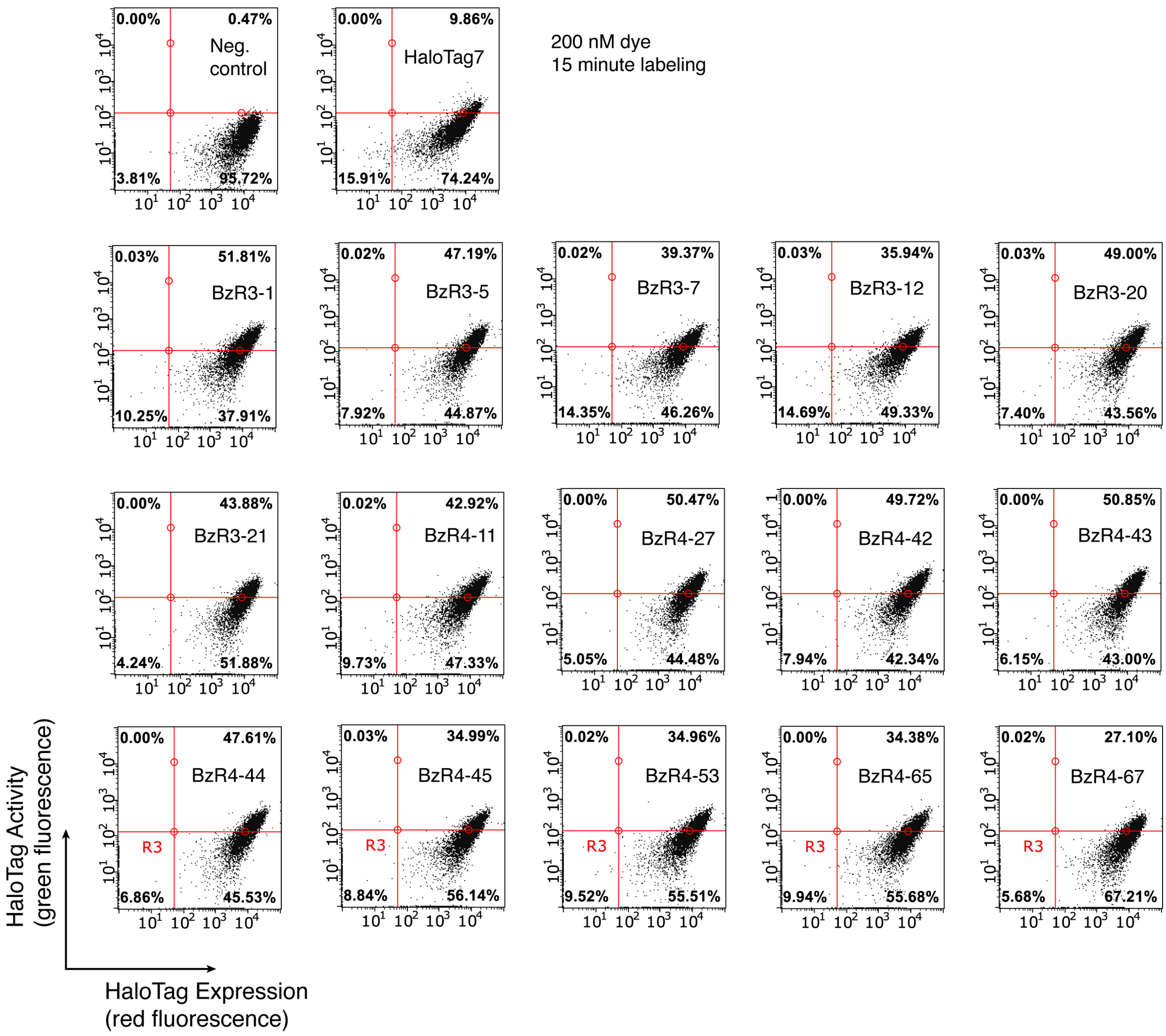

**Figure S5**. **Activity of screening hits, expressed on the surface of yeast analysis.** Single HaloTag variant clones were analyzed as described above. Cells were treated with 0.2 µM **Bz-1** for 15 minutes.

**Table S8. DNA sequences of selected HaloTag variants produced from yeast display experiments**.

| Name | Sequence |
|---|---|
| BzR3-1 (Variant 1) | ATGGCTGAAATTGGTACAGGTTTTCCATTTGATCCACATTACGTCGAAGTTT<br>TGGGTGAAAGAATGCATTACGTTGATGTTGGTCCAAGAGATGGTACACCA<br>GTTTTGTTTTTACATGGTAACCCAACTTCTTCATACGTTTGGAGAAACATCA<br>TCCCACATGTTGCGCCAACTCATAGATGTATTGCTCCAGATTTGATTGGTA<br>TGGGTAAATCTGATAAGCCAGATTTGGGTTATTTCTTTGATGATCATGTTAG<br>ATTCATGGATGCTTTTATTGAAGCATTGGGTTTGGAAGAAGTTGTTTTGGTT<br>ATTCATGATTGGGGTTCTGCATTAGGTTTTCATTGGGCTAAGAGAAACCCA<br>GAAAGAGTTAAGGGTATCGCTTTTATGGAGTTTATTAGACCAATTCCAACAT<br>GGGATGAATGGCCAGAATCTGCAAGAGAAACTTTCCAAGCTTTTAGAACAA<br>CTGATGTTGGTAGAAAGTTGATCATCGATCAAAACGTTTTTATTGAAGGTAC<br>ATTGCCAATGGGTGTTGTCAGACCATTGACTGAAGTTGAAATGGATCATTA<br>CAGAGAACCATTTTTAAACCCAGTTGATAGAGAACCATTGTGGAGATTTCC<br>AAATGAATTACCAATTGCTGGTGAACCAGCAAACATCGTTGCTTTGGTTGA<br>AGAATACATGGATTGGCTACATCAATCTCCAGTTCCAAAGTTGTTATTTTGG<br>GGTACACCAGGTGTTTTCATTCCACCAGCAGAAGCTGCAAGACTGGCTAA<br>GTCATTGCCAAACTGTAAGGCAGTTGATATTGGTCCAGGTTTGAATTTGTT<br>GCAAGAAGATAACCCAGATTTGATTGGTTCTGAAATTGCTAGATGGTTGTC<br>AACTTTA |
| BzR3-20 (Variant 3) | ATGGCTGAAATTGGTACAGGTTTTCCATTTGATCCACATTACGTTGAAGTTT<br>TGGGTGAAAGAATGCATTACGTTGATGTTGGTCCAAGAGATGGTACACCA<br>GTTTTGTTTTTACATGGTAACCCAACTTCTTCATACGTTTGGAGAAACATCA<br>TCCCACATGTTGCACCAACTCATAGATGTATTGCTCCAGATTTGATTGGTAT<br>GGGTAAATCTGATAAGCCAGATTTGGGTTATTTCTTTGATGATCATGTTAGA<br>TTCATGGATGCTTTTATTGAAGCATTGGGTTTGGAAGAAGTTGTTTTGGTTA<br>TTCATGATTGGGGTTCTGCATTAGGTTTTCATTGGGCTAAGAGAAACCCAG<br>AAAGAGTTAAGGGTATCGCTTTTATGGAGTTTATTAGACCAATTCCAACATG<br>GGATGAATGGCCAGAATTTGCAAGAGAAACTTTCCAAGCTTTTAGAACAAC<br>TGATGTTGGTAGAAAGTTGATCATCGATCAAAACGTTTTTATTGAAGGTACA<br>TTGCCAATGGGTGTTGTCAGACCATTGACTGAAGTTGAGATGGATCATTAC<br>AGAGAACCATTTTTAAGCCCAGTTGATAGAGAACCATTGTGGAGATTTCCA<br>AATGAATTACCAATTGCTGGTGAACCAGCAAACATCGTTGCTTTGGTTGAA<br>GAATACATGGATTGGTTACATCAATCTCCAGTTCCAAAGTTGTTATTTTGGG<br>GTACACCAGGTGCTTTAATTCCACCAGCAAAAGCTGCAAGATTGGCTAAGT<br>CATTGCCAAACTGTAAGGCAGTTGATATTGGTCCAGGTTTGAATTTGTTGC<br>AAGAAGATAACCCAGATTTGATTGGTTCTGAAATTGCTAGATGGTTGTCAA<br>CTTTA |
| BzR4-27 (Variant 4) | ATGGCTGAAATTGGTACAGGTTTTCCATTTGATCCACATTACGTTGAAGTTT<br>TGGGTGAAAGAATGCATTACGTTGATGTTGGTCCAAGAGATGGTACACCA<br>GTTTTGTTTTTACATGGTAACCCAACTTCTTCATACGTTTGGAGAAACATCA |

| | |
|---|---|
| | TCCCACATGTTGCACCAACTCATAGATGTATTGCTCCAGATCTGATTGGTA TGGGTAAATCTGATAAGCCAGATTTGGGTTATTTCTTTGATGATCATGTTAG ATTCATGGATGCTTTTATTGAAGCATTGGGTTTGGAAGAAGTTGTTTTGGTT ATTCATGATTGGGGTTCTGCATTAGGTTTTCATTGGGCTAAGAGAAACCCA GAAAGAGTTAAGGGTATCGCTTTTATGGAGTTTATTAGACCAATTCCAACAT GGGATGAATGGCCAGAATTTGCAAGAGAAACTTTCCAAGCTTTTAGAACAA CTGATGTTGGTAGAAAGTGGATCATCGATCAAAACGTTTTTATTGAAGGTA CATTGCCAATGGGTGTTGTCAGACCATTGACTGAAGTTGAAATGGATCATT ACAGAGAACCATTTTTAAACCCAGTTGATAGAGAACCATTGTGGAGATTTC CAAATGAATTACCAATTGCTGGTGAACCAGCAAACATCGTTGCTTTGGTTG AAGAATACATGGATTGGTTACATCAATCTCCAGTTCCAAAGTTGTTATTTTG GGGTACACCAGGTGCTTTAATTCCACCAGCAGAAGCTGCAAGATTGGCTA AGTCATTGCCAAACTGTAAGGCAGTTGATATTGGTCCAGGTTTGAATTTGT TGCAAGAAGATAACCCAGATTTGGTTGGTTCTGAAATTGCTAGATGGTTGT CAACTTTA |
| BzR4-42 (Variant 6) | ATGGCTGAAATTGGTACAGGTTTTCCATTTGATCCACATTACGTTGAAGTTT TGGGTGAAAGAATGCATTACGTTGATGTTGGTCCAAGAGATGGTACACCA GTTTTGTTTTTACATGGTAACCCAACTTCTTCATACGTTTGGAGAAACATCA TCCCACATGTTGCACCAACTCATAGATGTATTGCTCCAGATTTGATTGGTAT GGGTAAATCTGATAAGCCAGATTTGGGTTATTTCTTTGATGATCATGTTAGA TTCATGGATGCTTTTATTGAAGCATTGGGTTTGGAAGAAGTTGTTTTGGTTA TTCATGATTGGGGTTCTGCATTAGGTTTTCATTGGGCTAAGAGAAACCCAG AAAGAGTTAAGGGTATCGCTTTTATGGAGTTTATTAGACCAATTCCAACATG GGATGAATGGCCAGAACTTGCAAGAGAAACTTTCCAAGCTTTTAGAACAAC TGATGTTGGTAGAAAGTTGATCATCGATCAAAACGTTTTTATTGAAGGTACA TTGCCAATGGGTGTTGTCAGACCATTGACTGAAGTTGAAATGGATCATTAC AGAGAACCATTTTTAAACCCAGTTGATAGAGAACCATTGTGGAGATTTCCA AATGAATTACCAATTGCTGGTGAACCAGCAAACATCGTTGCTTCGGTTGAA GAATACATGGATTGGTTACATCAATCTCCAGTTCCAAAGTTGTTATTTTGGG GTACACCAGGTGCTTTAATTCCACCAGCAGAAGCTGCAAGATTGGCTAAGT CATTGCCAAACTGTAAGGCAGTTGATATTGGTCCAGGTTTGAATTTGTTGC AAGAAGATAACCCAGATTTGATTGGTTCTGAAATTGCTAGATGGTTGTCAA CTTTA |
| BzR4-43 (Variant 5) | ATGGCTGAAATTGGTACAGGTTTTCCATTTGATCCACATTACGTTGAAGTTT TGGGTGAAAGAATGCATTACGTTGATGTTGGTCCAAGAGATGGTACACCA GTTTTGTTTTTACATGGTAACCCAACTTCTTCATACGTTTGGAGAAACATCA TCCCACATGTTGCACCAACTCATAGATGTATTGCTCCAGATTTGATTGGTAT GGGTAAATCTGATAAGCCAGATTTGGGTTATTTCTTTGATGATCATGTTAGA TTCATGGATGCTTTTATTGAAGCATTGGGTTTGGAAGAAGTTGTTTTGGTTA TTCATGATTGGGGTTCTGCATTAGGTTTTCATTGGGCTAAGAGAAACCCAG AAAGAGTTAAGGGTATCGCTTTTATGGAGTTTATCAGACCAATTCCAACAT |

| | |
|---|---|
| | GGGATGAATGGCCAGAATTTGCAAGAGAAACTTTCCAAGCTTTTAGAACAA<br>CTGATGTTGGTAGAAAGTTGATCATCGATCAAAACGTTTTTATTGAAGGTAC<br>ATTGCCAATGGGTGTTGTCAGACCATTGACTGAAGTTGAAATGGATCATTA<br>CAGAGAACCATTTTTAAACCCAGTTGATAGAGAACCATTGTGGAGATTTCC<br>AAATGAATTACCAGTTGCTGGTGAACCAGCAAACATCGTTGCTTTGGTTGA<br>AGAATACATGGATTGGTTACATCAATCTCCAGTTCCAAAGTTGTTATTTTGG<br>GGTACACCAGGTGCTTTAATTCCACCAGCAGAAGCTGCAAGATTGGCTAA<br>GTCATTGCCAAACTGTAAGGCAGTTGATATTGGTCCAGGTTTGAATTTGTT<br>GCAAGAAGATAACCCAGATTTGATTGGTTCTGAAATTGCTAGATGGTTGTC<br>AACTTTA |
| BzR4-44<br>(Variant 2) | ATGGCTGAAATTGGTACAGGTTTTCCATTTGATCCACATTACGTTGAAGTTT<br>TGGGTGAAAGAATGCATTACGTTGATGTTGGTCCAAGAGACGGTACACCA<br>GTTTTGTTTTTACATGGTAACCCAACTTCTTCATACGTTTGGAGAAACATCA<br>TCCCACATGTTGCACCAACTCATAGATGTATTGCTCCAGATTTGATTGGTAT<br>GGGTAAATCTGATAAGCCAGATTTGGGTTATTTCTTTGATGATCATGTTAGA<br>TTCATGGATGCTTTTATTGAAGCATTGGGTTTGGAAGAAGTTGTTTTGGTTA<br>TTCATGATTGGGGTTCTGCATTAGGTTTTCATTGGGCTAAGAGAAACCCAG<br>AAAGAGTTAAGGGTATCGCTTTTATGGAGTTTATTAGACCAATTCCAACATG<br>GGATGAATGGCCAGAATTTGCAAGAGAAACTTTCCAAGCTTTTAGAACAAC<br>TGATGTTGGTAGAAAGTTGATCATCGATCAAAACGTCTTTATTGAAGGTACA<br>TTGCCAATGGGTGTTGTCAGACCATTGACTGAAGTTGAAATGGATCATTAC<br>AGAGAACCATTTTTAAACCCAGTTGATAGAGAACCATTGTGGAGATTTCCA<br>AATGAATTACCAATTGCTGGTGAACCAGCAAACATCGTTGCTTTGGTTGAA<br>GAATACATGGATTGGTTACATCAATCTCCAGTTCCAAAGTTGTTATTTTGGG<br>GTACACCAGGTGCTTTAATTCCACCAGCAGAAGCTGCAAGATTGGCTAAGT<br>CATTGCCAAACTGTAAGGCAGTTGATATTGGTCCAGGTTTGAATTTGTTGC<br>AAGAAGATAACCCAGATTTGATTGGTTCTGAAATTGCTAGATGGTTGTCAA<br>CTTTA |
| Variant 7 | ATGGCTGAAATTGGTACAGGTTTTCCATTTGATCCACATTACGTTGAAGTTT<br>TGGGTGAAAGAATGCATTACGTTGATGTTGGTCCAAGAGATGGTACACCA<br>GTTTTGTTTTTACATGGTAACCCAACTTCTTCATACGTTTGGAGAAACATCA<br>TCCCACATGTTGCACCAACTCATAGATGTATTGCTCCAGATTTGATTGGTAT<br>GGGTAAATCTGATAAGCCAGATTTGGGTTATTTCTTTGATGATCATGTTAGA<br>TTCATGGATGCTTTTATTGAAGCATTGGGTTTGGAAGAAGTTGTTTTGGTTA<br>TTCATGATTGGGGTTCTGCATTAGGTTTTCATTGGGCTAAGAGAAACCCAG<br>AAAGAGTTAAGGGTATCGCTTTTATGGAGTTTATTAGACCAATTCCAACATG<br>GGATGAATGGCCAGAACTTGCAAGAGAAACTTTCCAAGCTTTTAGAACAAC<br>TGATGTTGGTAGAAAGTTGATCATCGATCAAAACGTTTTTATTGAAGGTACA<br>TTGCCAATGGGTGTTGTCAGACCATTGACTGAAGTTGAAATGGATCATTAC<br>AGAGAACCATTTTTAAACCCAGTTGATAGAGAACCATTGTGGAGATTTCCA<br>AATGAATTACCAATTGCTGGTGAACCAGCAAACATCGTTGCTTCGGTTGAA |

| | |
|---|---|
| | GAATACATGGATTGGTTACATCAATCTCCAGTTCCAAAGTTGTTATTTTGGG GTACACCAGGTGCTTTTATTCCACCAGCAGAAGCTGCAAGATTGGCTAAGT CATTGCCAAACTGTAAGGCAGTTGATATTGGTCCAGGTTTGAATTTGTTGC AAGAAGATAACCCAGATTTGATTGGTTCTGAAATTGCTAGATGGTTGTCAA CTTTA |
| Variant 8 | ATGGCTGAAATTGGTACAGGTTTTCCATTTGATCCACATTACGTTGAAGTTT TGGGTGAAAGAATGCATTACGTTGATGTTGGTCCAAGAGATGGTACACCA GTTTTGTTTTTACATGGTAACCCAACTTCTTCATACGTTTGGAGAAACATCA TCCCACATGTTGCACCAACTCATAGATGTATTGCTCCAGATTTGATTGGTAT GGGTAAATCTGATAAGCCAGATTTGGGTTATTTCTTTGATGATCATGTTAGA TTCATGGATGCTTTTATTGAAGCATTGGGTTTGGAAGAAGTTGTTTTGGTTA TTCATGATTGGGGTTCTGCATTAGGTTTTCATTGGGCTAAGAGAAACCCAG AAAGAGTTAAGGGTATCGCTTTTATGGAGTTTATTAGACCAATTCCAACATG GGATGAATGGCCAGAACTGGCAAGAGAAACTTTCCAAGCTTTTAGAACAAC TGATGTTGGTAGAAAGTTGATCATCGATCAAAACGTTTTTATTGAAGGTACA TTGCCAATGGGTGTTGTCAGACCATTGACTGAAGTTGAAATGGATCATTAC AGAGAACCATTTTTAAACCCAGTTGATAGAGAACCATTGTGGAGATTTCCA AATGAATTACCAATTGCTGGTGAACCAGCAAACATCGTTGCTTCGGTTGAA GAATACATGGATTGGTTACATCAATCTCCAGTTCCAAAGTTGTTATTTTGGG GTACACCAGGTGCTTTAATTCCACCAGCAGAAGCTGCAAGATTGGCTAAGT CATTGCCAAACTGTAAGGCAGTTGATATTGGTCCAGGTTTGAATTTGTTGC AAGAAGATAACCCAGATTTGATTGGTTCTGAAATTGCTAGATGGTTGTCAA CTTTA |
| Variant 9 | ATGGCTGAAATTGGTACAGGTTTTCCATTTGATCCACATTACGTTGAAGTTT TGGGTGAAAGAATGCATTACGTTGATGTTGGTCCAAGAGATGGTACACCA GTTTTGTTTTTACATGGTAACCCAACTTCTTCATACGTTTGGAGAAACATCA TCCCACATGTTGCACCAACTCATAGATGTATTGCTCCAGATTTGATTGGTAT GGGTAAATCTGATAAGCCAGATTTGGGTTATTTCTTTGATGATCATGTTAGA TTCATGGATGCTTTTATTGAAGCATTGGGTTTGGAAGAAGTTGTTTTGGTTA TTCATGATTGGGGTTCTGCATTAGGTTTTCATTGGGCTAAGAGAAACCCAG AAAGAGTTAAGGGTATCGCTTTTATGGAGTTTATTAGACCAATTCCAACATG GGATGAATGGCCAGAACTGGCAAGAGAAACTTTCCAAGCTTTTAGAACAAC TGATGTTGGTAGAAAGTTGATCATCGATCAAAACGTTTTTATTGAAGGTACA TTGCCAATGGGTGTTGTCAGACCATTGACTGAAGTTGAAATGGATCATTAC AGAGAACCATTTTTAAACCCAGTTGATAGAGAACCATTGTGGAGATTTCCA AATGAATTACCAATTGCTGGTGAACCAGCAAACATCGTTGCTTCGGTTGAA GAATACATGGATTGGTTACATCAATCTCCAGTTCCAAAGTTGTTATTTTGGG GTACACCAGGTGCTTTTATTCCACCAGCAGAAGCTGCAAGATTGGCTAAGT CATTGCCAAACTGTAAGGCAGTTGATATTGGTCCAGGTTTGAATTTGTTGC AAGAAGATAACCCAGATTTGATTGGTTCTGAAATTGCTAGATGGTTGTCAA CTTTA |
| Variant 10 (BenzoHTag) | ATGGCTGAAATTGGTACAGGTTTTCCATTTGATCCACATTACGTTGAAGTTT TGGGTGAAAGAATGCATTACGTTGATGTTGGTCCAAGAGATGGTACACCA |

| | |
|---|---|
| | GTTTTGTTTTTACATGGTAACCCAACTTCTTCATACGTTTGGAGAAACATCA<br>TCCCACATGTTGCACCAACTCATAGATGTATTGCTCCAGATTTGATTGGTAT<br>GGGTAAATCTGATAAGCCAGATTTGGGTTATTTCTTTGATGATCATGTTAGA<br>TTCATGGATGCTTTTATTGAAGCATTGGGTTTGGAAGAAGTTGTTTTGGTTA<br>TTCATGATTGGGGTTCTGCATTAGGTTTTCATTGGGCTAAGAGAAACCCAG<br>AAAGAGTTAAGGGTATCGCTTTTATGGAGTTTATCAGACCAATTCCAACAT<br>GGGATGAATGGCCAGAATTTGCAAGAGAAACTTTCCAAGCTTTTAGAACAA<br>CTGATGTTGGTAGAAAGTTGATCATCGATCAAAACGTTTTTATTGAAGGTAC<br>ATTGCCAATGGGTGTTGTCAGACCATTGACTGAAGTTGAAATGGATCATTA<br>CAGAGAACCATTTTTAAACCCAGTTGATAGAGAACCATTGTGGAGATTTCC<br>AAATGAATTACCAGTGGCTGGTGAACCAGCAAACATCGTTGCTTTGGTTGA<br>AGAATACATGGATTGGTTACATCAATCTCCAGTTCCAAAGTTGTTATTTTGG<br>GGTACACCAGGTGCTTTAATTCCACCAGCAGAAGCTGCAAGATTGGCTAA<br>GTCATTGCCAAACTGTAAGGCAGTTGATATTGGTCCAGGTTTGAATTTGTT<br>GCAAGAAGATAACCCAGATTTGATTGGTTCTGAAATTGCTAGATGGTTGTC<br>AACTTTA |

**Table S9. Protein sequences of selected HaloTag variants produced from yeast display experiments.**

| Name | Sequence |
|---|---|
| BzR3-1<br>(Variant 1) | MAEIGTGFPFDPHYVEVLGERMHYVDVGPRDGTPVLFLHGNPTSSYVWRNIIP<br>HVAPTHRCIAPDLIGMGKSDKPDLGYFFDDHVRFMDAFIEALGLEEVVLVIHD<br>WGSALGFHWAKRNPERVKGIAFMEFIRPIPTWDEWPESARETFQAFRTTDVG<br>RKLIIDQNVFIEGTLPMGVVRPLTEVEMDHYREPFLNPVDREPLWRFPNELPIA<br>GEPANIVALVEEYMDWLHQSPVPKLLFWGTPGVFIPPAEAARLAKSLPNCKAV<br>DIGPGLNLLQEDNPDLIGSEIARWLSTL |
| BzR4-44<br>(Variant 2) | MAEIGTGFPFDPHYVEVLGERMHYVDVGPRDGTPVLFLHGNPTSSYVWRNIIP<br>HVAPTHRCIAPDLIGMGKSDKPDLGYFFDDHVRFMDAFIEALGLEEVVLVIHD<br>WGSALGFHWAKRNPERVKGIAFMEFIRPIPTWDEWPEFARETFQAFRTTDVG<br>RKLIIDQNVFIEGTLPMGVVRPLTEVEMDHYREPFLNPVDREPLWRFPNELPIA<br>GEPANIVALVEEYMDWLHQSPVPKLLFWGTPGALIPPAEAARLAKSLPNCKAV<br>DIGPGLNLLQEDNPDLIGSEIARWLSTL |
| BzR3-20<br>(Variant 3) | MAEIGTGFPFDPHYVEVLGERMHYVDVGPRDGTPVLFLHGNPTSSYVWRNIIP<br>HVAPTHRCIAPDLIGMGKSDKPDLGYFFDDHVRFMDAFIEALGLEEVVLVIHD<br>WGSALGFHWAKRNPERVKGIAFMEFIRPIPTWDEWPEFARETFQAFRTTDVG<br>RKLIIDQNVFIEGTLPMGVVRPLTEVEMDHYREPFLSPVDREPLWRFPNELPIA<br>GEPANIVALVEEYMDWLHQSPVPKLLFWGTPGALIPPAKAARLAKSLPNCKAV<br>DIGPGLNLLQEDNPDLIGSEIARWLSTL |
| BzR4-27<br>(Variant 4) | MAEIGTGFPFDPHYVEVLGERMHYVDVGPRDGTPVLFLHGNPTSSYVWRNIIP<br>HVAPTHRCIAPDLIGMGKSDKPDLGYFFDDHVRFMDAFIEALGLEEVVLVIHD<br>WGSALGFHWAKRNPERVKGIAFMEFIRPIPTWDEWPEFARETFQAFRTTDVG |

| | |
|---|---|
| | RKWIIDQNVFIEGTLPMGVVRPLTEVEMDHYREPFLNPVDREPLWRFPNELPI<br>AGEPANIVALVEEYMDWLHQSPVPKLLFWGTPGALIPPAEAARLAKSLPNCKA<br>VDIGPGLNLLQEDNPDLVGSEIARWLSTL |
| BzR4-43<br>(Variant 5) | MAEIGTGFPFDPHYVEVLGERMHYVDVGPRDGTPVLFLHGNPTSSYVWRNIIP<br>HVAPTHRCIAPDLIGMGKSDKPDLGYFFDDHVRFMDAFIEALGLEEVVLVIHD<br>WGSALGFHWAKRNPERVKGIAFMEFIRPIPTWDEWPEFARETFQAFRTTDVG<br>RKLIIDQNVFIEGTLPMGVVRPLTEVEMDHYREPFLNPVDREPLWRFPNELPV<br>AGEPANIVALVEEYMDWLHQSPVPKLLFWGTPGALIPPAEAARLAKSLPNCKA<br>VDIGPGLNLLQEDNPDLIGSEIARWLSTL |
| BzR4-42<br>(Variant 6) | MAEIGTGFPFDPHYVEVLGERMHYVDVGPRDGTPVLFLHGNPTSSYVWRNIIP<br>HVAPTHRCIAPDLIGMGKSDKPDLGYFFDDHVRFMDAFIEALGLEEVVLVIHD<br>WGSALGFHWAKRNPERVKGIAFMEFIRPIPTWDEWPELARETFQAFRTTDVG<br>RKLIIDQNVFIEGTLPMGVVRPLTEVEMDHYREPFLNPVDREPLWRFPNELPIA<br>GEPANIVASVEEYMDWLHQSPVPKLLFWGTPGALIPPAEAARLAKSLPNCKAV<br>DIGPGLNLLQEDNPDLIGSEIARWLSTL |
| (Variant 7) | MAEIGTGFPFDPHYVEVLGERMHYVDVGPRDGTPVLFLHGNPTSSYVWRNIIP<br>HVAPTHRCIAPDLIGMGKSDKPDLGYFFDDHVRFMDAFIEALGLEEVVLVIHD<br>WGSALGFHWAKRNPERVKGIAFMEFIRPIPTWDEWPELARETFQAFRTTDVG<br>RKLIIDQNVFIEGTLPMGVVRPLTEVEMDHYREPFLNPVDREPLWRFPNELPIA<br>GEPANIVASVEEYMDWLHQSPVPKLLFWGTPGAFIPPAEAARLAKSLPNCKAV<br>DIGPGLNLLQEDNPDLIGSEIARWLSTL |
| (Variant 8) | MAEIGTGFPFDPHYVEVLGERMHYVDVGPRDGTPVLFLHGNPTSSYVWRNIIP<br>HVAPTHRCIAPDLIGMGKSDKPDLGYFFDDHVRFMDAFIEALGLEEVVLVIHD<br>WGSALGFHWAKRNPERVKGIAFMEFIRPIPTWDEWPELARETFQAFRTTDVG<br>RKLIIDQNVFIEGTLPMGVVRPLTEVEMDHYREPFLNPVDREPLWRFPNELPV<br>AGEPANIVASVEEYMDWLHQSPVPKLLFWGTPGALIPPAEAARLAKSLPNCKA<br>VDIGPGLNLLQEDNPDLIGSEIARWLSTL |
| (Variant 9) | MAEIGTGFPFDPHYVEVLGERMHYVDVGPRDGTPVLFLHGNPTSSYVWRNIIP<br>HVAPTHRCIAPDLIGMGKSDKPDLGYFFDDHVRFMDAFIEALGLEEVVLVIHD<br>WGSALGFHWAKRNPERVKGIAFMEFIRPIPTWDEWPELARETFQAFRTTDVG<br>RKLIIDQNVFIEGTLPMGVVRPLTEVEMDHYREPFLNPVDREPLWRFPNELPV<br>AGEPANIVASVEEYMDWLHQSPVPKLLFWGTPGAFIPPAEAARLAKSLPNCKA<br>VDIGPGLNLLQEDNPDLIGSEIARWLSTL |
| (Variant 10)<br>**BenzoHTag** | MAEIGTGFPFDPHYVEVLGERMHYVDVGPRDGTPVLFLHGNPTSSYVWRNIIP<br>HVAPTHRCIAPDLIGMGKSDKPDLGYFFDDHVRFMDAFIEALGLEEVVLVIHD<br>WGSALGFHWAKRNPERVKGIAFMEFIRPIPTWDEWPELARETFQAFRTTDVG<br>RKLIIDQNVFIEGTLPMGVVRPLTEVEMDHYREPFLNPVDREPLWRFPNELPV<br>AGEPANIVALVEEYMDWLHQSPVPKLLFWGTPGALIPPAEAARLAKSLPNCKA<br>VDIGPGLNLLQEDNPDLIGSEIARWLSTL |

#### Recombinant Protein Expression

The six isolated DNA fragments and the yeast codon-optimized HaloTag7 sequence were amplified by PCR using primers **P10** and **P11** to introduce homologous overhangs and an N-terminal 6xHis sequence for cloning into the pET51(+)b plasmid between the NsiI and AvrII restriction sites. Cloned sequences were transformed into NEB BL21 competent *E. coli* by heat shock for protein expression and purification, as previously reported.[13] Recovered cells were grown on LB agar plates with ampicillin (100 μg/mL) overnight at 37 °C. Individual colonies were inoculated in 10 mL of LB broth with ampicillin and grown overnight at 37 °C. Overnight cultures were then diluted into 1 L of LB broth with ampicillin and grown for 2-4 hours at 37 °C until reaching an optical density at 600 nm of 0.6. Protein expression was induced with 0.5 mL IPTG (1 mM) for 4 hours at 37 °C or overnight at 20 °C before pelleting at 5,000 rpm for 10 min at 4 °C. Pellets were stored at -80 °C before lysis in sterile filtered His-binding buffer (50 mM Tris-Cl, 100 mM NaCl, 5 mM imidazole, 0.1 mM EDTA, 5 mM 2-mercaptoethanol, pH 8.0) with 30 mg lysozyme (Thermo Fisher Scientific) and ½ tablet cOmplete EDTA-free protease inhibitor (Roche). Lysed pellets were sonicated on ice for 10 minutes (10 seconds on/off, amplitude: 60%). Lysate was then incubated with Pierce Universal Nuclease for Cell Lysis (Thermo Fisher Scientific) for 5 minutes on ice and clarified by centrifugation for 30 minutes at 10,000 rpm at 4 °C before decanting and centrifuging again. The clarified lysate was purified by batch affinity purification with 3 mL HisPur™ Ni-NTA resin (Thermo Scientific). Before purification, Ni-NTA resin was washed and then equilibrated with His-binding buffer for >45 min on a rocking platform at 4 °C. After equilibration, clarified lysate was incubated with the Ni-NTA resin in His-binding buffer for 1 hour on a rocking platform at 4 °C. After draining the flowthrough, the Ni-NTA resin was washed 10 times: 3 times with His-binding buffer and 7 times with His-wash buffer (50 mM Tris-Cl, 500 mM NaCl, 10 mM imidazole, 0.1 mM EDTA, 5 mM 2-mercaptoethanol, pH 6.8). We found that washing with a higher salt concentration (500 mM NaCl rather than to 100 mM) and a lower pH (pH 6.8 rather than pH = 8.0) yielded purer HaloTag. Purified HaloTag was eluted from the resin with His-elution buffer (50 mM Tris-Cl, 50 mM NaCl, 300 mM imidazole, 0.1 mM EDTA, 5 mM 2-mercaptoethanol). Finally, eluted fractions were pooled and buffer exchanged into PBS using 7k MWCO Zeba Spin Desalting Columns (Thermo Fisher Scientific) before aliquoting and snap freezing in liquid nitrogen. Frozen protein samples were stored at -80 °C. This protocol routinely produced purified protein yields of ~20 mg/L with the yeast codon-optimized HaloTag sequences. By contrast, a bacteria codon optimized sequence produced ~90 mg/L in our hands.[13]

Variants 7-10 were generated using site-directed mutagenesis using Variant 5 and Variant 6 constructs as templates. Mutagenesis was done using the Q5® Site-Directed Mutagenesis Kit from NEB while following manufacturers procedures. Primers P14 and P15 were used to introduce F144L, P16 and P17 were used to introduce I211V, and P18 and P19 were used to introduce L246F. Successful mutagenesis was verified by Sangar sequencing of ten random colonies and variants were expressed and purified following the same protocol from above.

#### Kinetics

Conjugation rates of **Bz-1** and related substrates to HaloTag7 and identified variants were measured as previously described using a fluorescence plate reader.[14] Briefly, 0.5 μM substrate in PBS + 0.1% Tween was added to varying concentrations of 2x protein (4, 3, 2, 1.5, 1, 0.5 μM) in PBS + 0.1% Tween in a 1:1 ratio. Final dye concentration was 0.25 μM and final protein concentrations were 2, 1.5, 1, 0.75, 0.5, and 0.25 μM. Fluorescence (excitation 450 nm, emission 535 nm) was measured as a function of time accounting for the gap between addition of protein and the first measurement (<20 seconds). Fluorescence was measured every 15 seconds for 30 minutes. Kinetic traces were normalized and fit to a one-phase association exponential curve (Equation 1) where $Y_0$ was set to 0. All results reported are the average of at least three independent biological replicates and errors are reported as the standard error of the mean (Table S10, Fig. S6 & S7). We note that the larger error associated with the faster measured Variant 7 and 10 are a result of these rates approaching the limit of detection using traditional plate reader fluorescence measurements. Stopped-flow kinetics will be more suitable for rates any faster than $10^6$ $M^{-1}s^{-1}$.

$$(1)\ Y = Y_0 + (Plateau - Y_0)\left(1 - e^{-k_{obs}X}\right)$$

Conjugation rates of CA-TMR with benzoHTag and select variants were measured by fluorescent plate reader and fluorescence polarization as previously reported.[13] Rates between 0.05 μM substrate and identical protein concentrations as above were used. Conjugation rates of CA-$JF_{635}$ were measured by monitoring the growth of fluorescence emission at 665 nm. As noted in the main text, larger concentration of dye (500 μM), and thus higher protein concentrations (100, 50, 25, 12.5, 6.25, and 3.125 μM) were used to detect a large enough change in fluorescence upon reacting with benzoHTag.

**Table S10. Second-order rate constants for Bz-1 Conjugation to HaloTag7, and Variants 1-6.**

| | $k_{app}$ ($M^{-1}s^{-1}$) | | | | | |
|---|---|---|---|---|---|---|
| | Trial 1 | Trial 2 | Trial 3 | Trial 4 | Trial 5 | Average |
| HaloTag7 | 1229 | 1944 | 1541 | 1194 | - | 1477 ± 174 |
| Variant 1 | 6264 | 5501 | 9910 | 8152 | - | 9329 ± 2023 |
| Variant 2 | 13088 | 9708 | 11804 | 9430 | - | 11008 ± 873 |
| Variant 3 | 5335 | 8117 | 16818 | 11482 | 7648 | 8317 ± 997 |
| Variant 4 | 13849 | 10632 | 9803 | 7098 | - | 10346 ± 1390 |
| Variant 5 | 41090 | 43023 | 36382 | 39734 | 46113 | 41268 ± 1625 |
| Variant 6 | 33688 | 31757 | 28303 | 40014 | - | 33441 ± 2458 |
| Variant 7 | 56574 | 14313 | 77788 | 36241 | 59796 | 68844 ± 8161 |
| | 62359 | 78259 | 104179 | 131318 | 96792 | |
| | 40027 | 50245 | 95765 | 35912 | 93088 | |
| Variant 8 | 12232 | 12721 | 16085 | 19501 | 17559 | 15620 ± 1395 |
| Variant 9 | 34860 | 19916 | 51566 | 46157 | 48808 | 44601 ± 4149 |
| | 48551 | 53187 | 36241 | 62127 | | |
| Variant 10 | 61758 | 147628 | 58690 | 102928 | 75640 | 95366 ± 9249 |
| **BenzoHTag** | 89957 | 110609 | 109900 | 101182 | | |

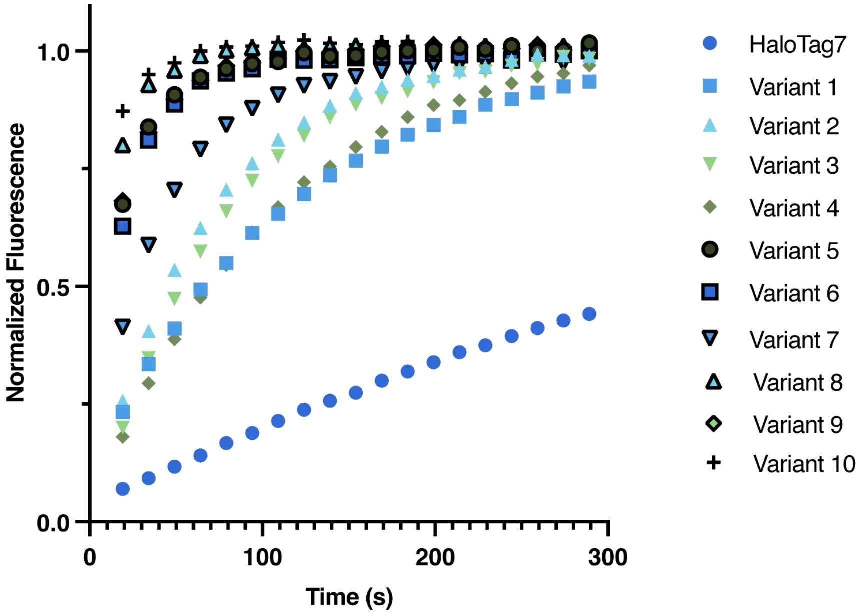

**Figure S6. Representative kinetic traces Bz-1 conjugation to HaloTag7, BenzoHTag, and Varians 1-9.** Example kinetic traces of 0.25 µM **Bz-1** reacting with 1.0 µM of protein for the first five minutes of measurement.

a

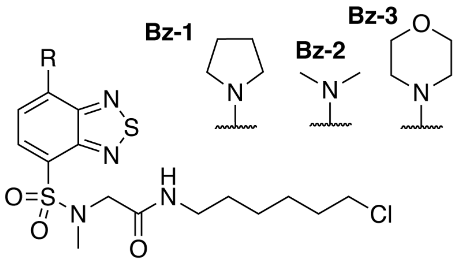

b

| | $k_{app}$ ($M^{-1}s^{-1}$) (x$10^3$) | | |
|---|---|---|---|
| | HaloTag7 | Variant 6 **BenzoHTag** | Variant 5 |
| **Bz-1** | 1.47 ± 0.17 | 33.44 ± 2.4 | 40.06 ± 1.40 |
| **Bz-2** | 0.32 ± 0.00 | 3.72 ± 0.12 | 4.04 ± 0.44 |
| **Bz-3** | 0.24 ± 0.01 | 2.97 ± 0.30 | 2.57 ± 0.21 |

c

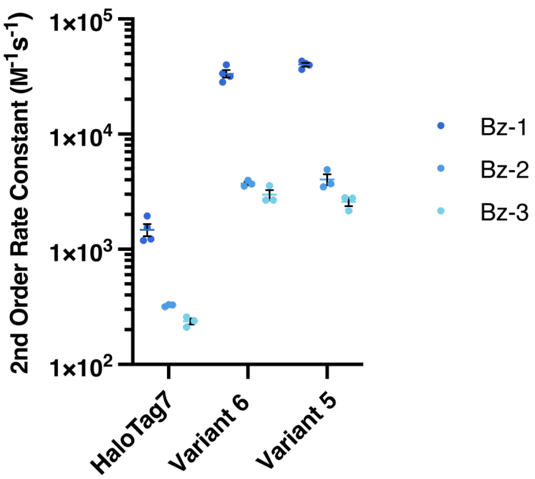

**Figure S7. Reaction rates of other benzothiadiazole substrates with HaloTag7, BenzoHTag, and Variant 5.** (a) Structures of **Bz-1, Bz-2,** and **Bz-3**. (b,c) Summary of second-order rate constants of **Bz-1, Bz-2,** and **Bz-3** with HaloTag7, BenzoHTag, and Variant 5. All results reported are the average of at least three independent biological replicates and errors are reported as the standard error of the mean.

**Table S11. Second-order rate constants for CA-TMR and CA-JF$_{635}$ Conjugation to BenzoHTag and select variants.**

| | $k_{app}$ ($M^{-1}s^{-1}$) | | | | | |
|---|---|---|---|---|---|---|
| | Trial 1 | Trial 2 | Trial 3 | Trial 4 | Trial 5 | |
| | **CA-TMR** | | | | | Average |
| **BenzoHTag** | 16975 | 22330 | 21751 | 22986 | 16975 | 21011 ± 1369 |
| Variant 6 | 42499 | 45729 | 53562 | - | - | 47263 ± 3284 |
| Variant 8 | 17186 | 17390 | 18765 | - | - | 17780 ± 495.8 |
| Variant 9 | 232 | 2160 | 506 | - | - | 966.0 ± 602.2 |
| | **CA-JF$_{635}$** | | | | | Average |
| **BenzoHTag** | 79.54 | 132.4 | 129 | - | - | 113.6 ± 17.08 |

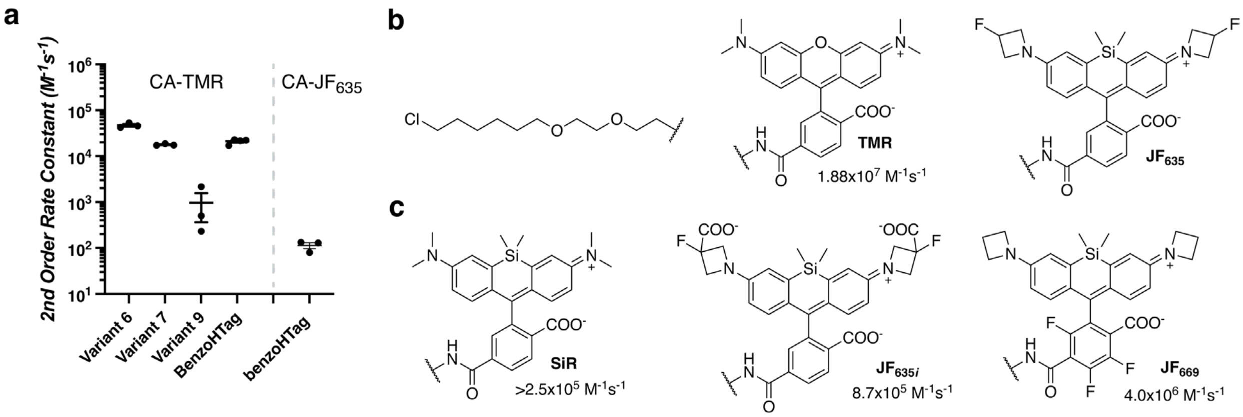

**Figure S8. Reaction rates of rhodamine chloroalkane substrates with HaloTag7, BenzoHTag, and Variant 6.** (a) Plot of apparent second order rate constants of **CA-TMR** and **CA-JF$_{635}$** with BenzoHTag and selected variants. (b) Structures of **CA-TMR** and **CA-JF$_{635}$**. (c) Structures of analogous silicon rhodamine dyes, **SiR**[15]; **JF$_{635i}$**[16]; **JF$_{669}$**[17, 18], and their apparent second order rate constants used to estimate the conjugation rate of **CA-JF$_{635}$** to HaloTag7.

#### Photophysical measurements

Steady state photophysical measurements were conducted in aerated solvents at approximately 10 μM of dye such that the absorbance did not exceed 0.1. Quantum yield of fluorescence was calculated using eq. 1, where F is the integrated intensities, *f* is the overlap absorbance value between the dye and standard, and η is the refractive index of the solvent; *i* and s stand for sample and standard, respectively. Coumarin153 in ethanol was used as the standard ($\varphi_{Fl,\ standard}$ = 0.53). Properties were measured in duplicate.

$$(\mathbf{1})\ \Phi_{fl}^{i} = \left(\frac{F^{i} f_{s} \eta_{i}^{2}}{F^{s} f_{i} \eta_{s}^{2}}\right) \Phi_{fl}^{s}$$

| | $\Phi_{fl}$ | $\varepsilon$ ($M^{-1}cm^{-1}$) | $\lambda_{abs,\ max}$ (nm) | $\lambda_{em,\ max}$ (nm) |
|---|---|---|---|---|
| BenzoHTag•**Bz-1** | 0.58 | 8100 | 446 | 530 |

**Mammalian Cell Culture**

*General* – HaloTag7 and BenzoHTag were subjected to PCR using primers **P12** and **P13** to introduce homologous overhangs for cloning into pCDNA/FRT/TO_TOMM20 plasmid (Gift from Kai Johnsson,[5] pCDNA5/FRT/TO_TOMM20_HaloTag9 was a gift from Kai Johnsson (Addgene plasmid # 169335) between the AgeI and XhoI restriction sites.[5] After cloning, these plasmids were transformed into NEB 5α competent *E. coli* and plasmid isolated by maxi-prep (Invitrogen).

U-2 OS cells (ATCC) were cultured in high glucose MyCoy's 5A media (Thermo Fisher Scientific) supplemented with 10% fetal bovine serum and grown at 37 °C in a humidified environment of 5% $CO_2$. Two days prior to an experiment, 10,000 cells were seeded per well in a 96-well plate and grown for 24 hours. Cells were then transiently transfected with expression plasmid using X-tremeGene 9 transfection reagent (Roche) according to manufacturer protocols. Cells were incubated with transfection mix for 18-24 hours.

*Flow Cytometry* – Transfection media was aspirated, and cells were washed with PBS. Non-transfected cells and cells transfected with either HaloTag7 or BenzoHTag, and non-transfected cells were then treated with varying concentrations of **Bz-1** or **Bz-3** 100 µL in Opti-MEM reduced serum media (Thermo Fisher Scientific) for 10 minutes. Media was then aspirated, and cells were trypsinized without wash steps from the well using 0.05% Trypsin in PBS, diluted to 200 µL. 10,000 live, single cells were analyzed by flow cytometry. The **Bz-1** fluorescence (green) of each cell was measured using a 488 nm laser and a 525 nm emission filter, and the **CA-JF$_{635}$** fluorescence (red) was measured using a 642 nm laser and a 661 nm emission filter. The mean green fluorescence of 5,000 cells was calculated and the background due to cell autofluorescence was subtracted using measurement from non-transfected, non-dye treated cells. Signal-over-background was calculated by taking the ratio of green fluorescence between dye-treated, transfected cells and dye-treated, non-transfected of cells. The reported mean green fluorescence values are for the entire analyzed population and are thus likely and underestimation of the true enhancement of green fluorescence in live cells as it includes ~80% of cells that were not transfected.

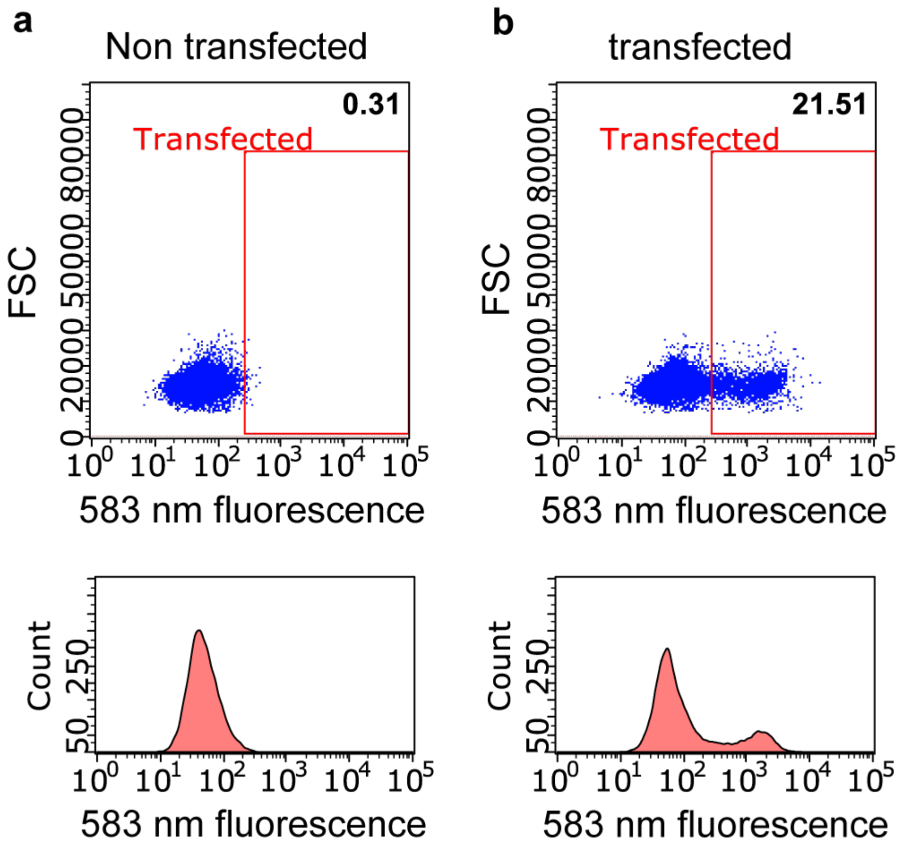

**Figure S9. Transient transfection efficiencies for live cell experiments.** Representative examples of raw flow cytometry data indicating transfection efficiencies. To verify and quantitate transfection, cells were treated with excess (1.0 µM) CA-TMR for 15 minutes, followed by a wash-out step and then analyzed by flow cytometry using the Blue (488 nm) excitation source and Yellow (583 nm) emission filter. The gate was made relative to cells that were no transfected but treated with dye.

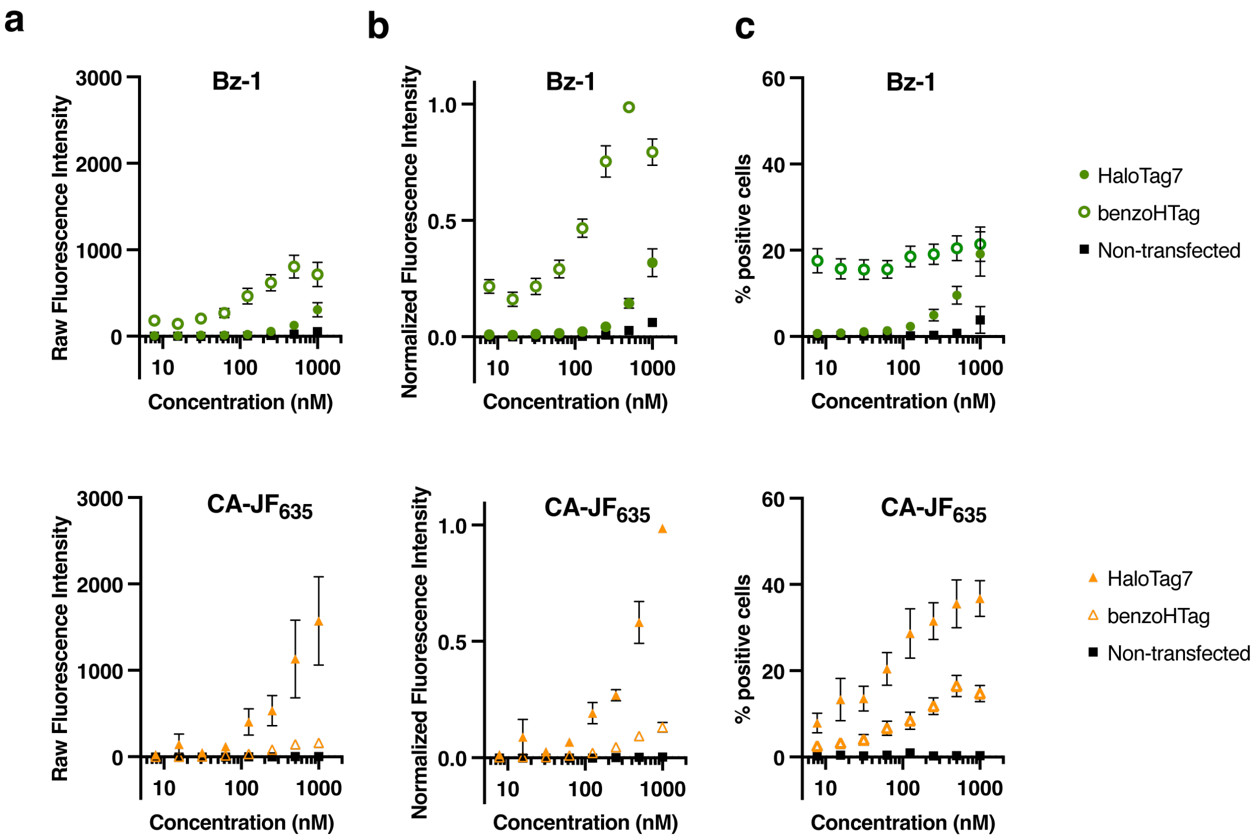

**Figure S10. Flow-cytometry data of BenzoHTag or HaloTag expressing cells.** (a) Dose-response raw fluorescence intensity of **Bz-1** or **CA-JF**$_{635}$ treated U-2 OS cells or U-2 OS cells transiently expressing H2B fusions fo HaloTag7 or BenzoHTag. Cells were treated for 10 minutes prior to analysis. (b) Normalized fluorescence intensity. (c) Dose-response of the percentage of **Bz-1** or **CA-JF**$_{635}$ labeled U-2 OS cells, or U-2 OS cells expressing HaloTag7 or BenzoHTag above background autofluorescence of untreated control cells.

*Confocal Microscopy* – Transfection media was aspirated, and cells were treated with Hoescht stain (1 μg/mL) for 30 minutes in Opti-MEM. Cells were washed twice with PBS and then treated with Opti-MEM for an extended 30-minute wash step. 50 μL of HEPES-buffered saline (HBS) were added to the cells. Prior to imaging, cells were then treated with 50 μL of 250 nM or 20 nM of **Bz-1** in HBS (final concentration 125 nM or 10 nM) and imaged directly without removing excess dye. Images were acquired on a Leica TCS SP8 microscope and processed using ImageJ software.

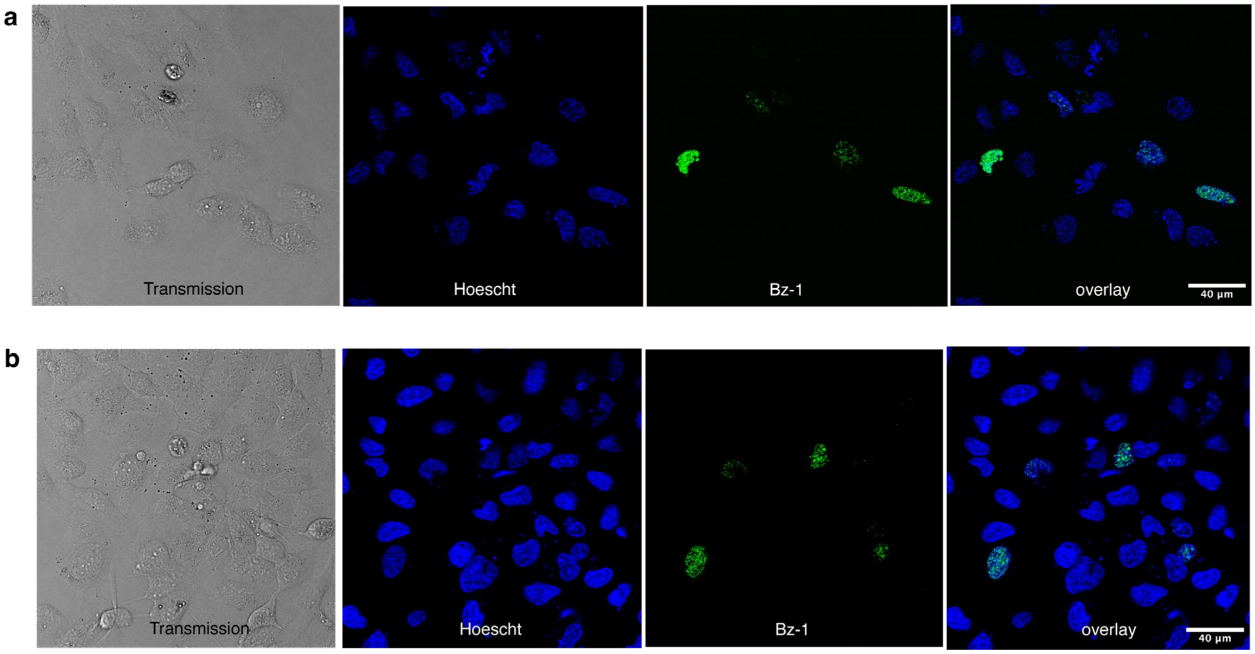

**Figure S11. Turn-on fluorescence in BenzoHTag-expressing cells.** Confocal microscopy images of **Bz-1** with U-2 OS cells that were transiently transfected with a H2B-BenzoHTag fusion, treated with (a) 10 nM of **Bz-1** or (b) 125 nM of **Bz-1** and imaged directly without washing. Blue is Hoescht staining, and green fluorescence is **Bz-1**.

*Live cell labeling kinetics* – Transfected cells were labeled with Hoescht stain as described above. To each well, 50 μL HBS was added. Shortly after the start of video recording, 50 μL of **Bz-1** in HBS was added to a final concentration of 125 or 10 nM. The appearance green fluorescence of transfected cells was recorded in real-time. Videos were recorded at 6 frames/second for five minutes. Videos were analyzed using ImageJ with the Time Series Analyzer V3 plugin.[1] The fluorescence intensity of at least 10 transfected cells (as determined by the last frame) was measured every 25 frames (~4 seconds) and plotted as a function of time. The average background fluorescence of 10 non-transfected cells was also measured every 25 frames and was subtracted from the fluorescence intensity of the transfected cells. Averages of the ≥10 transfected cells are shown in Fig. 4c in the main text. Curves were fit for each kinetic trace for each of the ≥10 analyzed transfected cells and measured $k_{app}$ ($s^{-1}$) are shown in Figure S12.

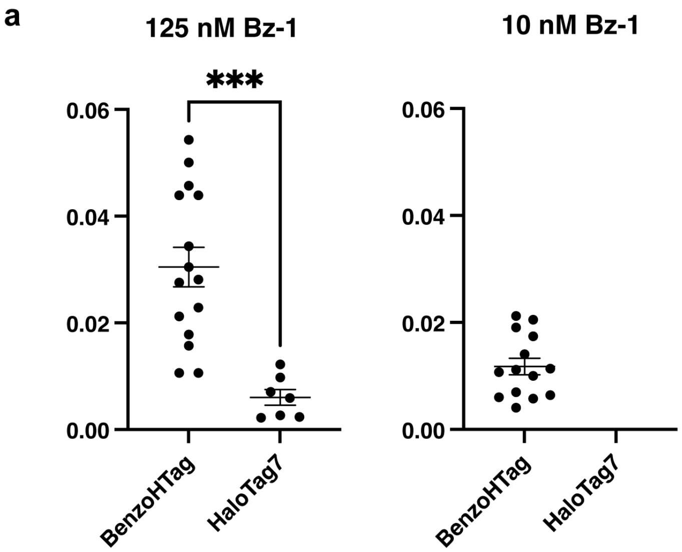

b

| | $k_{obs}$ ($s^{-1}$) | |
|---|---|---|
| | 125 nM | 10 nM |
| BenzoHTag | 0.03 ± 0.003 | 0.012 ± 0.001 |
| HaloTag7 | 0.006 ± 0.001 | ND |

**Figure S12. Summary of live-cell labeling kinetics of U-2 OS cells transiently transfected with BenzoHTag or HaloTag and treated with Bz-1.** Rates were calculated by fitting curves to each of the ≥10 transfected cells analyzed. Significance was determined by an unpaired Student's t-test; p=0.0003. (a) Labeling rates for cells treated with 125 nM (left) or 10 nM (right) **Bz-1**. (b) Summary of average $k_{obs}$ values ($s^{-1}$). Errors are standard error of the mean.

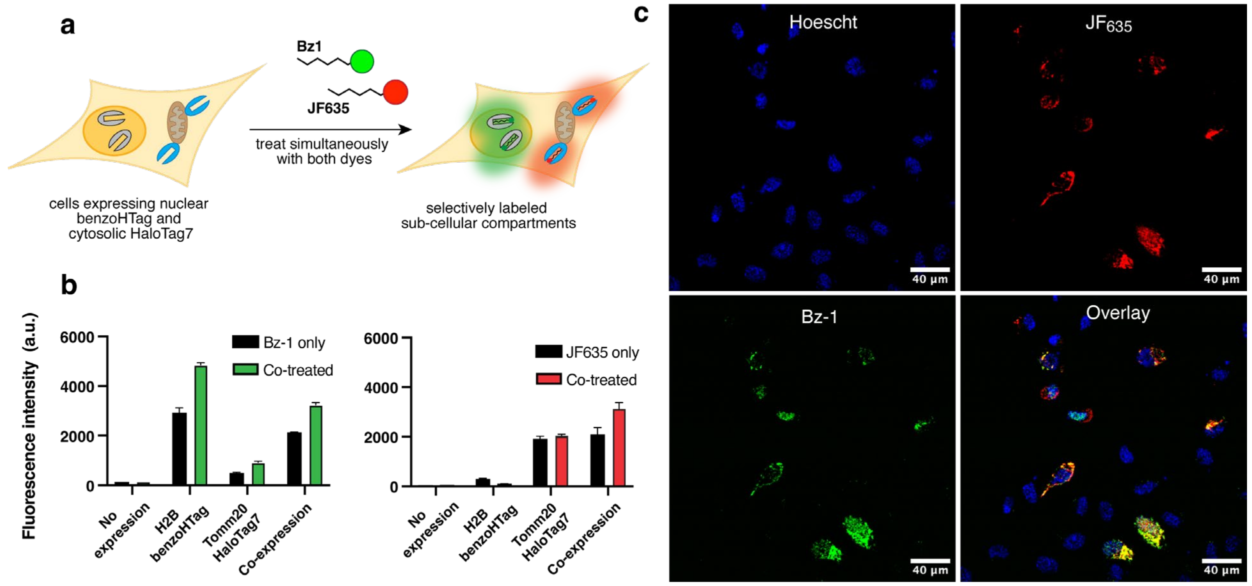

**Figure S13. Simultaneous multiplexed labeling with Bz-1 and CA-JF$_{635}$.** (a) U-2 OS cells co-expressing H2B-BenzoHTag and Tomm20-HaloTag7 were labeled simultaneously with a solution of

125 nM of each **Bz-1** and **CA-JF$_{635}$** and analyzed without washing out excess dye. (b) Flow cytometry data of the brightest 15% of cells in orthogonal labeling experiments with 125 nM **Bz-1** and 125 nM of **CA- JF$_{635}$** for 10 minutes. Raw flow cytometry histograms can be found in Fig. S14. U-2 OS cells expressing H2B-benzoHTag, Tomm20-HaloTag7, or with side-by-side comparison of cells treated with only dye or a co-treated with both dyes. (c) Confocal microscopy images of U-2 OS cells transiently transfected with both H2B-BenzoHTag and Tomm20-HaloTag7. Cells were treated with 125 nM **Bz-1** and 125 nM **CA- JF$_{635}$** for 60 minutes and are co-stained with nuclear Hoescht dye.

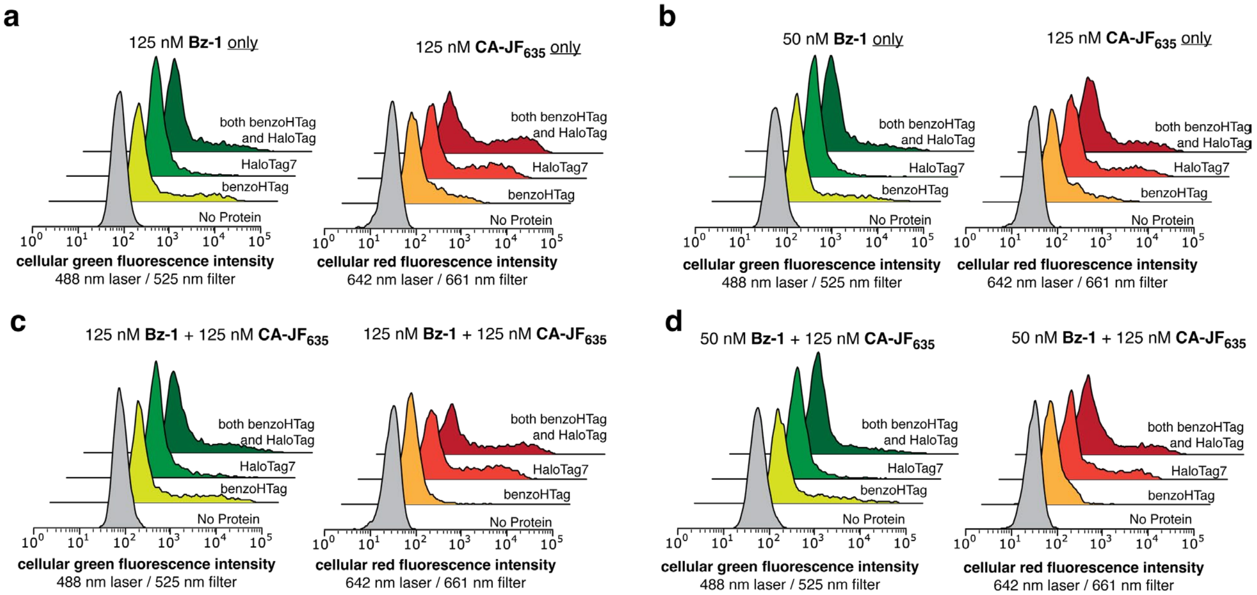

**Figure S14. Histogram of simultaneous multiplexed labeling of cells co-expressing HaloTag7 and BenzoHTag.** U-2 OS that were co-transfected with Tomm20-HaloTag7 and H2B-BenzoHTag constructs were treated either with a solution of only **Bz-1**, **CA-JF$_{635}$,** or a mixture of both dyes, and analyzed by flow cytometry. (a, c) Cells treated with 125 nM of **Bz-1,** 125 nM of **CA-JF$_{635}$**, or a solution of both dyes for 10 minutes. (b,d) Cells treated with 50 nM of **Bz-1,** 125 nM of **CA-JF$_{635}$**, or a solution both dyes for 60 minutes. The fluorescence intensity of the top 15% of cells were extracted for analysis.

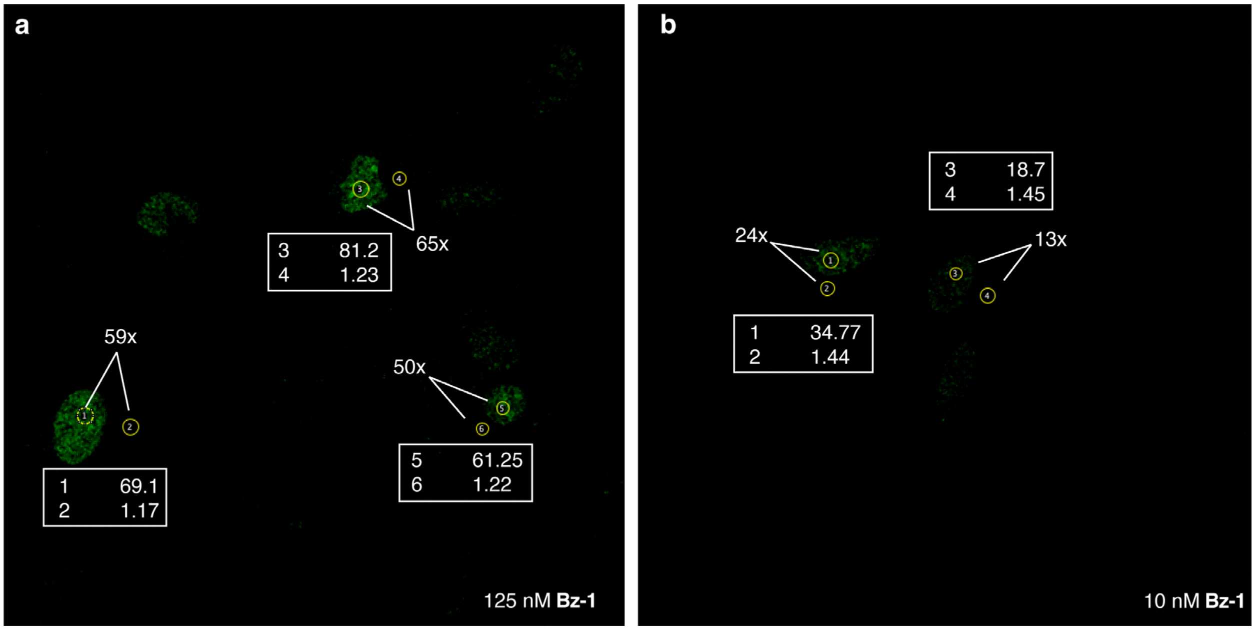

**Figure S15. Signal over background of Bz1•benzoHTag in live cell imaging.** Signal over background in live U2OS cells expressing benzoHTag as a Tomm20 fusion treated with (a) 125nM or (b) 10 nM of **Bz1** for ~60 minutes and imaged without washing or replacing the media. Mean fluorescence intensity of signal (nuclear fluorescence) over background (cytosolic fluorescence) are indicated. Analysis was conducted using ImageJ on raw image files obtained from a confocal fluorescence microscope.

**References**

1. Schneider, C. A.; Rasband, W. S.; Eliceiri, K. W., NIH Image to ImageJ: 25 years of image analysis. *Nature Methods* **2012,** *9* (7), 671-675.
2. Los, G. V., et al., HaloTag: A Novel Protein Labeling Technology for Cell Imaging and Protein Analysis. *ACS Chemical Biology* **2008,** *3* (6), 373-382.
3. Encell, L. P., et al., Development of a dehalogenase-based protein fusion tag capable of rapid, selective and covalent attachment to customizable ligands. *Curr Chem Genomics* **2012,** *6*, 55-71.
4. Erdmann, R. S., et al., Labeling Strategies Matter for Super-Resolution Microscopy: A Comparison between HaloTags and SNAP-tags. *Cell Chemical Biology* **2019,** *26* (4), 584-592.e6.
5. Frei, M. S., et al., Engineered HaloTag variants for fluorescence lifetime multiplexing. *Nature Methods* **2021**.
6. Cook, A.; Walterspiel, F.; Deo, C., HaloTag-Based Reporters for Fluorescence Imaging and Biosensing. *ChemBioChem* **2023,** *24* (12), e202300022.
7. Chao, G., et al., Isolating and engineering human antibodies using yeast surface display. *Nat Protoc* **2006,** *1* (2), 755-68.

8. Islam, M., et al., Chemical Diversification of Simple Synthetic Antibodies. *ACS Chemical Biology* **2021,** *16* (2), 344-359.
9. Angelini, A., et al., Protein Engineering and Selection Using Yeast Surface Display. Springer New York: 2015; pp 3-36.
10. Van Deventer, J. A.; Wittrup, K. D., Yeast Surface Display for Antibody Isolation: Library Construction, Library Screening, and Affinity Maturation. Humana Press: 2014; pp 151-181.
11. Branon, T. C., et al., Efficient proximity labeling in living cells and organisms with TurboID. *Nat Biotechnol* **2018,** *36* (9), 880-887.
12. Colby, D. W., et al., Engineering antibody affinity by yeast surface display. *Methods Enzymol* **2004,** *388*, 348-58.
13. Deprey, K.; Kritzer, J. A., HaloTag Forms an Intramolecular Disulfide. *Bioconjugate Chemistry* **2021,** *32* (5), 964-970.
14. Liu, Y., et al., The Cation−π Interaction Enables a Halo-Tag Fluorogenic Probe for Fast No-Wash Live Cell Imaging and Gel-Free Protein Quantification. *Biochemistry* **2017,** *56* (11), 1585-1595.
15. Lukinavičius, G., et al., A near-infrared fluorophore for live-cell super-resolution microscopy of cellular proteins. *Nature Chemistry* **2013,** *5* (2), 132-139.
16. Jonker, C. T. H., et al., Accurate measurement of fast endocytic recycling kinetics in real time. *Journal of Cell Science* **2020,** *133* (2).
17. Grimm, J. B., et al., A general method to optimize and functionalize red-shifted rhodamine dyes. *Nature Methods* **2020,** *17* (8), 815-821.
18. Wilhelm, J., et al., Kinetic and Structural Characterization of the Self-Labeling Protein Tags HaloTag7, SNAP-tag, and CLIP-tag. *Biochemistry* **2021**.
